# Multi-omic analyses reveal a differential contribution of chromatin-associated PP1 holoenzymes to mitotic exit and G1 re-establishment

**DOI:** 10.1101/2024.04.29.591626

**Authors:** Konstantinos Stamatiou, Florentin Huguet, Marta Budzinska, Ines J deCastro, Christos Spanos, Juri Rappsilber, Paola Vagnarelli

## Abstract

Mitotic exit is an important part of the cell cycle that requires coordination of many chromatin and cytoskeleton remodelling events to successfully complete cell division and maintain cell identity. Protein de-phosphorylation is a key step in directing mitotic exit and Protein phosphatase (PP1) is essential to this process, however the specific contribution of its numerous targeting subunits is still unknown. Here we have investigated the function of three chromatin-associated PP1 targeting subunits in exiting mitosis Repo-Man, Ki-67 and PNUTS. We have generated endogenously tagged, auxin-degradable alleles for each subunit and used a multi-omic approach to address their specific contribution towards transcription resumption, chromatin accessibility and protein phosphorylation at the transition from mitosis to G1. This approach has identified their distinct role in mitotic exit, provided unique datasets for the cell cycle community, and highlighted novel functions for Ki-67 and Repo-Man in genome stability and organisation.

## INTRODUCTION

When cells exit mitosis, a complex but coordinated series of morphological and biochemical changes need to occur in order to re-establish a functional G1 nucleus that is either ready for the next cell cycle or drives differentiation. At this cell cycle transition, the mitotic spindle is disassembled, the mitotic chromosomes are reorganised into the interphase chromatin, the kinetochores are dismantled and transcription resumes. In recent years, several studies have investigated some of these aspects and we start gaining a clearer picture of its regulation.

Upon mitotic exit, mitotic chromatin reorganises into the interphase form where topologically associated domains (TADs) are re-established (Abramo et al., 2019), and the chromatin entanglements present on mitotic chromosomes are quickly removed, driven by chromatin expansion, with remaining entanglements being then resolved during early G1; this is mediated by Topo II and failure in the process delays the re-establishment of heterochromatic and euchromatic domains (Hildebrand et al., 2024).

These changes are also associated with the repositioning of chromosomes in their interphase configuration but the molecular mechanisms driving it are not known. A-type lamins (and in some cells the Lamin B receptor -LBR) seem to play a role as they are essential for tethering the heterochromatin at the nuclear envelope (Solovei et al., 2013). More recently, it was shown that Lamin C, but not Lamin A, is required for the 3D organization of Lamin Associated domains (LADs) and overall chromosome organization (Wong et al., 2021) where Lamin A S22 phosphorylation appears to be mediating the process (Wong *et al*., 2021).

The transition into G1 is also accompanied by waves of transcription reactivation where the amplitude of transcription at a specific subset of genes is highly increased (Palozola et al., 2017). This could be explained either by the re-establishment of the long-range chromatin interactions that would bring enhancers close to their promoters and/or by increased binding or sequential de-phosphorylation of transcription factors.

Indeed, protein de-phosphorylation is one the main mechanisms driving mitotic exit. In mammalian cells, Protein Phosphatase 1 (PP1) and Protein Phosphatase 2A (PP2A) orchestrate and drive the direction of mitotic exit events. In fact, even in the absence of APC/C activity or protein translation, dephosphorylation still occurs after CDK1 inhibition (Cundell et al., 2016) (Wu et al., 2009). Dephosphorylation rates correlate with the sequence characteristics of each substrate and define the specificity of the individual phosphatases (Cundell *et al*., 2016).

Nevertheless, we are still far from understanding the contribution of each holoenzyme in these processes. PP1 acts as a multimeric complex composed of a catalytic subunit and one or two regulatory/targeting subunits (RIPPOs), where the latter are essential to direct the localisation of the phosphatase activity. In this respect, considering the major changes occurring during miotic exit at a chromatin level in terms of structure, positioning and transcriptional activity, understanding the role of chromatin-associated phosphatases is becoming a priority in the field.

RNAi based approaches have identified Repo-Man (CDCA2) (Trinkle-Mulcahy et al., 2006), Ki-67 (Booth et al., 2014) and PNUTS (Kim et al., 2003) as important chromatin-associated PP1 regulatory subunits. Repo-Man has been shown to dephosphorylate Histone H3 (T3, S10 and S28) (Qian et al., 2011) (de Castro et al., 2017; Vagnarelli et al., 2011) and Lamin A S22 during mitotic exit (Huguet et al., 2022) (Moriuchi and Hirose, 2021) thus playing a role in re-establishing heterochromatin (de Castro *et al*., 2017); Ki-67 has been linked to the regulation of heterochromatin (Sobecki et al., 2016) (Sun et al., 2017) and pericentromeric chromatin (van Schaik et al., 2022), the nucleolar positioning of the NOR containing chromosomes (Booth *et al*., 2014) and Xi heterochromatin maintenance (Sun *et al*., 2017); PNUTS has been shown to be an important negative regulator of RNA pol II elongation rate (Landsverk et al., 2020). The suggested function of PNUTS also includes modulation of tumor suppressor genes, such as retinoblastoma (Rb) and sequestering PTEN (Udho et al., 2002) (Krucher et al., 2006) (Kavela et al., 2013). Moreover, PNUTS enhances in vitro chromatin decondensation in a PP1-dependent manner (Landsverk et al., 2005).

However, because the majority of these studies have been conducted by RNAi based approaches, it is not possible to rule out that the phenotypes observed are consequences of defects originated by a role that these proteins play in other stages of the cell cycle. We therefore decided to evaluate the direct contribution of chromatin-associated PP1 holocomplexes in establishing the G1 nucleus by using three HCT116 cell lines where we have tagged the endogenous Repo-Man or Ki-67 or PNUTS with the AID module. Addition of auxin allows the degradation of the protein within 4 h and the consequences of a mitotic exit in the absence of this protein can be directly evaluated. Using these cell lines together with a multi-omic approach, we have revealed that each holoenzyme has a very specific contribution to the G1 establishment. Repo-Man and Ki-67 degradation leads to a strong arrest at the restriction checkpoint while PNUTs degradation causes a major de-regulation of transcription. Moreover, we have unveiled an important function for Repo-Man in sustaining the spindle assembly checkpoint and a novel role for Ki-67 in maintaining centromere integrity in mitosis. This study represents the first major advance towards understanding the function and specificity of PP1 homocomplexes at the M/G1 transition, it provides unique and precious datasets for these holocomplexes and sets the parameters for analysing cell cycle transition stages.

## Results

### Differential contribution of Repo-Man, Ki-67 and PNUTS to the establishment of the transcription landscape in G1

To assess the specific contribution of 3 major chromatin-associated PP1 targeting subunits to the establishment of the G1 nucleus, we needed to discriminate between their roles at this specific stage of the cell cycle form the secondary effects caused by their functions prior to mitosis. To this purpose, we used a Ki-67 endogenously tagged with the AID degron module (Stamatiou et al., 2024) and we generated two new similar cell lines in HCT116 where either the endogenous alleles of Repo-Man or PNUTS were tagged at the C-terminus with the mClover:AID module (Supplementary Figure 1 A, B, E, F). The new cell lines were genotyped by PCR (Supplementary Figure 1 C, G, H) and the tagged alleles verified by western blotting (Supplementary Figure 1 D, I). In all these cell lines, OSTR1 is expressed under a doxycycline (Dox) inducible promoter and the addition of Dox and Auxin (IAA) leads to degradation of the proteins within 3-4 hours (Figure 1 B-D and Supplementary Figure 1 D, I).

**Figure 1.**
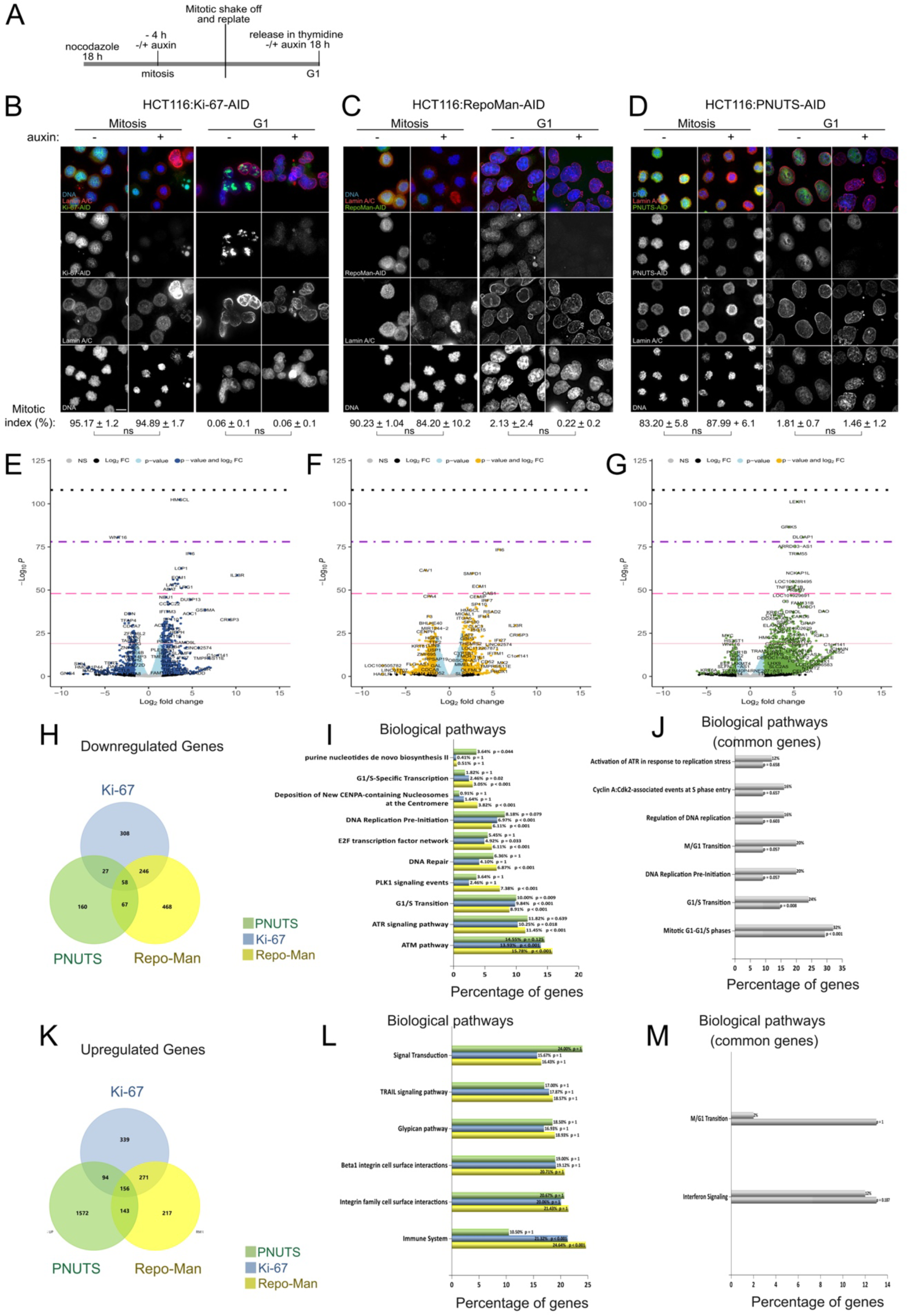
A) Scheme of the experiment: after 18 h in nocodazole, cells were treated with or without IAA for 4 h, followed by mitotic shake-off and cells replated and released in Thymidine with or without IAA for 18 h (G1). B, C and D) Representative images of LaminA/C immunostaining using anti-LaminA/C antibody on HCT116:Ki-67-AID (B), HCT116:RepoMan-AID (C), and HCT116:PNUTS-AID (D) cell lines in mitosis (left panel) and G1 (right panel) without or with IAA. Scale bar 5 μm. Mitotic index analyses were performed using 3 biological replicates for each cell line, the values represent the average of 3 independent replicas and ± Standard deviations. Samples size: HCT116:Ki-67-AID (B); Mitosis: Control=1695, IAA=1601 and G1: Control=1610, IAA=1705, HCT116:RepoMan-AID (C); Mitosis: Control=703, IAA=954 and G1: Control=1191, IAA=1203, and HCT116:PNUTS-AID (D); Mitosis: Control=677, IAA=643 and G1: Control=775, IAA=847. The experiments were analysed by Chi-squared test. ns= not significant. E, F and G) Volcano plots of the differentially expressed genes identified by RNA-seq for the HCT116:Ki-67-AID (E), HCT116:RepoMan-AID (F), and HCT116:PNUTS-AID (G) cell lines in G1 following the experiment in (A). Pink line p-value < 10e-20, Hot pink line p-value=10e-20-10e-30, purple line p-value=10e-20-10e-60 and black line p-value=10e-20-10e-90. H and K) Venn diagrams of the downregulated (H) and upregulated (K) genes identified by RNA-seq of the cell lines in E, F and G. I and L) FunRich biological pathways enrichment analysis of the downregulated (I) and upregulated (L) genes identified by RNA-seq of the cell lines in E, F and G. J and M) FunRich biological pathways enrichment analysis of the downregulated common (I) and upregulated common (L) genes between the cell lines, identified by RNA-seq of the cell lines in E,F and G.

Using these new cell lines, we were in the position of addressing the question about their specific contributions at the M/G1 transition and G1 chromatin landscape establishment.

The experimental setup for the study was chosen based on previous reports describing either the transcription resumption (Palozola *et al*., 2017) or the de-phosphorylation kinetics (Holder et al., 2020) after mitosis.

We therefore arrested the cells with Nocodazole in presence of DOX for 18 h, then cells were treated with or without IAA for 4 h; mitotic cells were collected by mitotic shake off and then released from the prometaphase block in normal medium with thymidine (with or without IAA to harvest all the cells at the G1/S boundary (Figure 1 A). In these experimental conditions, all the cell lines blocked well in prometaphase upon Nocodazole treatment (Figure 1 B, C, D – Mitosis and Mitotic Index), and the degradation of each protein did not affect the release from the block, as no mitotic cells were observed at the collection time point (Figure 1 B, C, D – G1 and Mitotic Index). As expected, degradation of Ki-67 in mitosis causes the well described phenotype of chromosome clustering (Cuylen-Haering et al., 2020) (suggesting that its function is not only required during chromosome condensation but also continuously during mitosis (Figure 1 B - Mitosis, + IAA). No obvious phenotypes in terms of chromosome morphology were observed upon degradation of PNUTS or Repo-Man at this stage (Figure 1 C, D - Mitosis, + IAA). Upon release, all the cells exited mitosis, chromosomes decondensed and the nuclear envelope (at least by Lamin A/C staining) reformed. By all these parameters we conclude that, without each PP1 holoenzyme, cells can progress from mitosis to G1.

One of the key transitions necessary to set up a G1 nuclear landscape is the ordered resumption of transcription (Palozola *et al*., 2017). We therefore tested how transcription resumption was affected by a mitotic exit in the absence of the 3 RIPPOs.

We performed RNA sequencing using cells arrested at the G1/S boundary following the protocol in Figure 1 A. The comparison between the three cell lines without IAA showed very little differences among them suggesting that the tags on the different genes have little influence on this stage of the cell cycle (Supplementary Figure 2 A). However, the comparison with the IAA treated cells showed a very different and protein-specific transcription landscape. In fact, only 58 downregulated genes were common between the 3 protein degradations and the highest number of shared downregulated genes were between Repo-Man and Ki-67 while PNUTS did not presented many down-regulated genes (Figure 1 E-H). Moreover, the majority of differentially expressed genes were specific for each protein: this shows that gene-specificity was prominent above a general G1 nuclear disorganisation response. While Ki-67 and Repo-Man have roughly a similar number of genes that were down or up-regulated upon a mitotic exit in the absence of the protein, PNUTS behaved very differently with a very high proportion of genes that were up-regulated compared to the ones downregulated when cells entered G1 in the absence of the protein (Figure 1 K). This already classifies these RIPPOs in two distinct categories. The data so far indicate that PNUTS does not affect chromatin decondensation as previously suggested (Landsverk *et al*., 2005) but also that it is essential for a balanced transcription resumption after mitosis.

We then further analysed the biological pathways of the dysregulated genes and their relative transcription factors to evaluate if specific features could be assigned to a specific protein. For the downregulation, G1/S transcription, DNA replication pre-initiation, E2F transcription factor network, ATM and ATR signalling pathway were significantly downregulated in the absence of both Ki-67 and Repo-Man (Figure 1 I); PLK1 and DNA repair were only affected by lack of Repo-Man and the Purine de novo biosynthesis II was only affected by lack of PNUTS (Figure 1 I). Among the common downregulated genes only the G1/S pathway was significantly affected thus suggesting that somehow all the proteins pay a role (although unique) to set up the G1 nucleus and, if missing, the cell cycle transition to the S phase is halted (Figure 1 J). However, Repo-Man and Ki-67 share downregulated genes that belong not only to G1/S checkpoint but also to the activation of the pre-replicative complex, and activation of the ATR pathway in response to replicative stress (Supplementary Figure 2 J). The degradation of each protein causes a significant decrease in genes transcribed by the E2F1 transcription factor but Repo-Man and PNUTS also include genes transcribed by NFYA and SP1 respectively (Supplementary Figure 2 B-D).

No specific pathways were found upregulated in these experiments apart from the Immune system for Repo-Man and Ki-67 (Figure 1 L, M, and Supplementary Figure 1 K), with a subset of genes shared by all three RIPPOs but not significantly enriched (Figure 1 M).

All these analyses strongly suggest that the 3 RIPPOs are essential for a proper re-organisation of the G1 nucleus and, in their absence, S phase cannot occur as most of the E2F1 targets are not expressed; nevertheless, they differ in their specificity.

### Lack of Repo-Man and Ki-67 during mitotic exit triggers a G1 arrest at the restriction checkpoint and activation of the interferon response

The transcriptomic data suggested that all the proteins were somehow altering the transcription landscape with E2F1 gene targets being downregulated. This scenario would well fit with a G1 checkpoint activation (Fischer et al., 2022).

To further analyse the transcriptomic in a timeline context and identify the stage where the transcription alteration occurs, we have cross referenced our new datasets with the transcription resumption after mitosis timeline obtained by Palozola et al. (Palozola *et al*., 2017). Although the experiments were conducted in a different cell line, their data suggested that the cell-line specific genes are activated at later timepoints, making housekeeping and cell cycle genes still a valid comparison. The intersect of their datasets with the differentially expressed genes in our transcriptomics, shows that mitotic exit in the absence of each RIPPO affects mainly the resumption of transcription of genes activated 80 min after release (in the Palozola et al. experiment set up timeline) (Figure 2 A – 80’ dataset). Genes that do not switch on at 80 min upon a mitotic exit in the absence of Repo-Man or Ki-67 belong to the category of G1/S transition, DNA damage and DNA replication (Figure 2 B).

**Figure 2.**
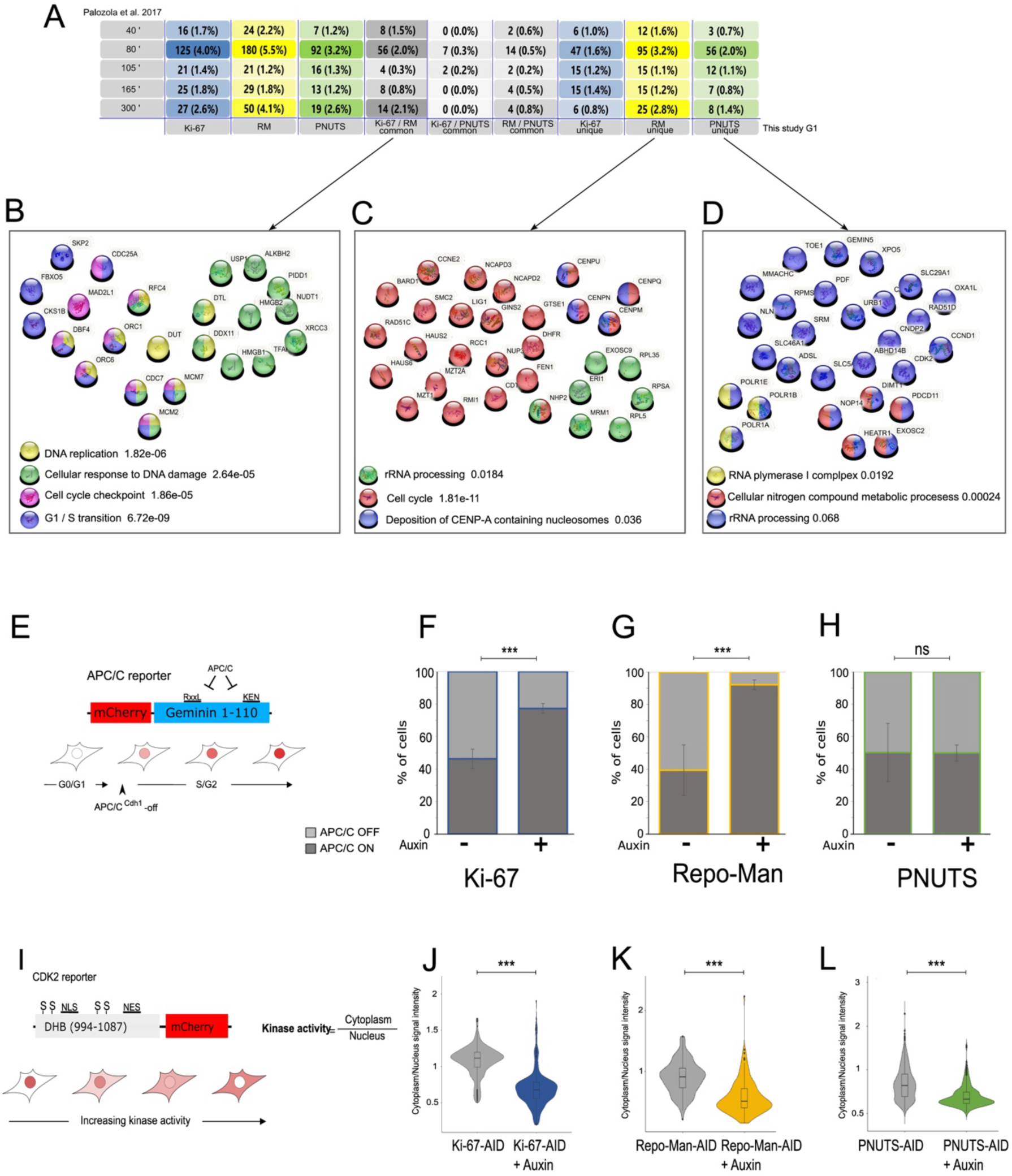
A) Venn diagram of the (Palozola *et al*., 2017) datasets (the number indicates the time points in their experimental set-up) and the downregulated genes of the HCT116:Ki-67-AID (Ki-67), HCT116:RepoMan-AID (RM) and HCT116:PNUTS-AID (PNUTS) cell lines, downregulated common genes between HCT116:Ki-67-AID and HCT116:RepoMan-AID genes (Ki-67 / RM common), downregulated common genes between HCT116:Ki-67-AID and HCT116:PNUTS-AID genes Ki-67 / PNUTS common), downregulated common genes between HCT116:RepoMan-AID and HCT116:PNUTS-AID genes (RM / PNUTS common), downregulated unique genes of HCT116:Ki-67-AID (Ki-67 unique), downregulated unique genes of HCT116:RepoMan-AID (RM unique) and downregulated unique genes of HCT116:PNUTS-AID (PNUTS unique). B) String analysis of overlapping common downregulated genes between HCT116:Ki-67-AID, HCT116:RepoMan-AID and 80 min time point of (Palozola *et al*., 2017). C) String analysis of overlapping unique downregulated genes of HCT116:RepoMan-AID and 80 min time point of (Palozola *et al*., 2017). D) String analysis of overlapping unique downregulated genes of HCT116:PNUTS-AID and 80 min time point of (Palozola *et al*., 2017). E) Schematic illustration of the APC/CCDH1 reporter (Spencer *et al*., 2013) used for the generation of HCT116:Ki-67-AID Geminin, HCT116:RepoMan-AID Geminin and HCT116:PNUTS-AID Geminin cell lines for the experiments in F,G and H, respectively. F, G and H) The graphs represent the percentage of Geminin positive and negative cells of HCT116:Ki-67-AID (F), HCT116:RepoMan-AID (G) and HCT116:PNUTS-AID (H) cell lines in the experiment as it described in Figure 1 A. The values are the average of 3 biological replicas and the error bars represent the standard deviations. Samples size: HCT116:Ki-67-AID : Control=1197, IAA=1685, HCT116:RepoMan-AID: Control=699, IAA=832 and HCT116:PNUTS-AID: Control=1174, IAA=709. The experiments were analysed by Chi-squared test. ***= p< 0.001 and ns= not significant. I) Schematic illustration of CDK2 reporter (Spencer *et al*., 2013), used for the generation of HCT116:Ki-67 AID DHB, HCT116:RepoMan-AID DHB and HCT116:PNUTS-AID DHB cell lines for the experiments in J, K and L J, K and L) The violin plots represent the distribution of the cytoplasmic/nuclear signals of 3 biological replicates of the HCT116:Ki-67-AID DHB (J), HCT116:RepoMan-AID DHB (K) and HCT116:PNUTS-AID DHB (L) in the experiment as described in Figure 1A. The box inside the violins represents the 75th and 25th percentile, whiskers the upper and lower adjacent values and the line is the median. Samples size: HCT116:Ki-67-AID: Control=329, IAA=310, HCT116:RepoMan-AID: Control=164, IAA=174 and HCT116:PNUTS-AID: Control=550, IAA=462. The data were analysed with a Wilcoxon test. ***=p<0.001.

Lack of Ki-67 does not block the re-activation of a specific pathway; genes that fail to increase transcription at this stage, specifically upon Repo-Man degradation, include rRNA processing, CENP-A deposition, and cell cycle (Figure 2 C). Interestingly, Repo-Man is normally upregulated at this stage of the cell cycle as shown in (Palozola *et al*., 2017) dataset and this could suggest that this holoenzyme represents a key regulator of this cell cycle transition, a function so far unknown. Lack of PNUTS during mitotic exit affects the upregulation of RNA polymerase I complex and rRNA processing (Figure 2 D).

These analyses indicate that resumption of transcription at 80 min is affected when mitotic exit occurs without these PP1 holoenzymes and again provides a picture of a very protein-specific contribution to the G1 transcription landscape.

Nevertheless, the major block in transcription resumption and the E2F1 target genes downregulation strongly point towards a checkpoint activation (Fischer *et al*., 2022) caused by the lack of these proteins in G1.

In G1, cells need to make decisions between entering the cell cycle (Restriction checkpoint) and then committing to duplicate the DNA. The first step is marked by the inactivation of the APC/C and then the activation of CDK2. To address if mitotic exit in the absence of these PP1 targeting subunits would affect this progression, we generated new cell lines on the AID background with reporters for either APC/C^CDH1^ (Figure 2 E) or CDK2 activity (Figure 2 I) (Spencer et al., 2013). Inactivation of the APC/C^CDH1^ would cause the mCherry reporter to accumulate in the nucleus. The analyses of cells that went through a mitotic exit without Repo-Man or Ki-67 showed that these cells cannot inactivate the APC/C^CDH1^ (cells are mCherry negative) and therefore are arrested at the restriction checkpoint (Figure 2 F, G). On the contrary, cells that exited mitosis without PNUTS can pass through this checkpoint stage even if the transcription landscape is aberrant (Figure 2 H). For the CDK2 reporter, activation of CDK2 kinase leads to the accumulation of the mCherry reporter in the cytoplasm. In this case, mitotic exit in the absence of each protein led to a block in CDK2 activation (Figure 2 I-L). These data show that all these RIPPOs contribute to the progression to G1 but the role they play is specific with PNUTS negative cells arresting later in G1. Interestingly, mis-regulation of transcription, as caused by lack of PNUTS, allows cells to pass the restriction checkpoint but it is detected by the G1/S transition checkpoint.

We then asked why lack of Repo-Man and Ki-67 would trigger a G1 block. The activation of the interferon pathway shown by the transcriptome analyses led us to consider that maybe genome instability and chromosome mis-segregation could be the major cause for the arrest. In fact, chromosome mis-segregation and micronuclei have been linked to the activation of the interferon response via the cGAS/STING pathway (Mackenzie et al., 2017).

We therefore analysed the ploidy of the interphase cells that exited mitosis with and without each of the RIPPOs. We conducted FISH on cells arrested at the G1/S boundary with two centromeric probes for chromosome 15 and chromosome 1. These analyses revealed that lack of Repo-Man and Ki-67 leads to a significant increase of aneuploid cells for both chromosome 1 and chromosome 15, however, lack of PNUTS did not (Supplementary Figure 3 C-E). These results are interesting because lack of PNUTS also triggered an interferon response but with less genes involved (Figure 1 M and Supplementary Figure 3 A, B) and no aneuploidy. In any case, some form of DNA damage should be present to cause the interferon activation, therefore we set up to analyse the presence of DNA damage using a comet assay. Cells treated as before (Figure 1 A) were subjected to the neutral COMET assay; the length of the comet tails that exited mitosis without each protein (+ IAA) was measured and compared to the ones of cells that completed mitotic exit in the presence of the proteins (- IAA). The analyses showed that lack of Repo-Man and Ki-67 cause a significant increase in aneuploidy and present DNA damage (Supplementary Figure 3 H-J). Interestingly, also lack of PNUTS causes an increase in DNA damage (Supplementary Figure 3 H-J) but not aneuploidy; for PNUTS the interferon response is weaker (less genes involved) and possibly not sufficiently strong to halt progression through the restriction checkpoint. These data could indicate a gene signature for a response to aneuploidy and a different one to DNA damage. Future work would investigate these relationships.

Another interesting and perhaps important observation that emerged from the transcriptome analyses is related to cells that are arrested in G1 upon Ki-67 degradation. We have already shown that cells without Ki-67 block in G1 for 2 days but then they overcome the block and can survive without Ki-67(Stamatiou *et al*., 2024); previous work has shown that cells lacking Ki-67 do not metastasize (Mrouj et al., 2021) (Zheng et al., 2009) (Kausch et al., 2004; Kausch et al., 2005). Here we have detected a significant enrichment for the biological pathway related to the mesenchymal-to-epithelial transition when cells exit mitosis without Ki-67 (Supplementary Figure 3 K). This observation is quite intriguing as Ki-67 is a well-known proliferation marker used for staging and prognosis of several cancer types; the finding that its degradation during mitotic exit leads to a reversal of the aggressive phenotype of cancer cells could be a useful and important step towards considering this marker not only as a bystander of proliferation but also as a cancer target.

### Chromatin accessibility and epigenetics marks only partially correlate

All these proteins have been linked in the past to chromatin organisation and their absence could potentially cause chromatin alterations that, in turn, contribute to the changes in transcription observed in G1. We have therefore analysed the chromatin status of the G1 nuclei by ATAC sequencing and by immunoblotting using several antibodies against histone modifications. The protocol used was the same as in the previous experiments (Figure 1 A). ATAC sequencing analyses revealed that some changes occurred within the chromatin, but their scale is much less than the ones observed by RNA sequencing. Lack of Ki-67 or Repo-Man during mitotic exit caused mainly an increase in regions that are less accessible (273 and 203 respectively) (Figure 3 A and B); on the contrary, PNUTS presented similar levels of changes in both directions (138 regions were less accessible and 149 more open) (Figure 3 C); again the level of changes is inferior to the scale of dysregulation of transcription described before. We also evaluated if the chromatin that changed accessibility upon degradation of these proteins was confined to specific regions of the genome. For Repo-Man and PNUTS the changes show a similar distribution where 84% fall within promoters (less than 1 Kb), distal intragenic and introns (Figure 3 D, F); however, for Ki-67, 79% of changes are located within the proximal promoters (less than 1 Kb) (Figure 3 D). Moreover, the regions that become less accessible upon degradation of these proteins are enriched for the open chromatin marks H3K9ac and H3K27ac and the same histone marks are also significantly enriched in regions that become more accessible upon PNUTS degradation (Figure 3 G, H).

**Figure 3.**
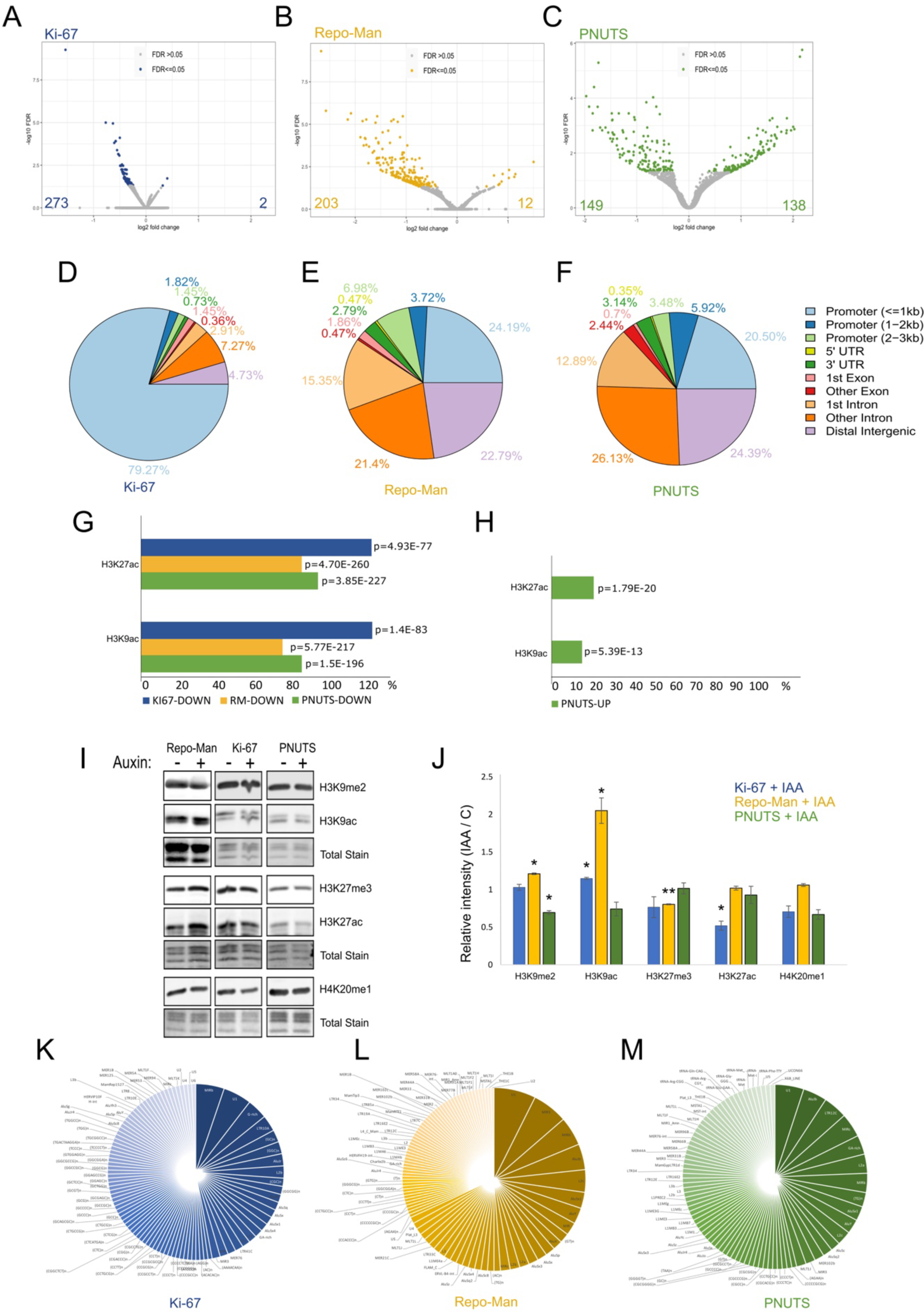
A, B and C) Volcano plots of the differentially accessible chromatin peaks obtained by ATAC-seq for the HCT116:Ki-67-AID (A), HCT116:RepoMan-AID (B), and HCT116:PNUTS-AID (C) cell lines following the experiment indicated in Figure 1 A. D, E and F) Genome-wide distribution of differentially open chromatin peaks obtained by ATAC-seq for the HCT116:Ki-67-AID (D), HCT116:RepoMan-AID (E), and HCT116:PNUTS-AID (F) cell lines following the experiment indicated in Figure 1 A. The accessible regions were divided into promoter (<= 1kb, 1-2kb and 2-3kb), gene body (5′UTR, 3′UTR, 1st exon, other exon, 1st intron and other intron), and distal intergenic regions. G) Overlaps between less accessible chromatin regions obtained by ATAC-seq for the HCT116:Ki-67-AID, HCT116:RepoMan-AID, and HCT116:PNUTS-AID cell lines following the experiment indicated in Figure 1 A, and HCT116 H3K9ac and H3K27ac ChIP-seq datasets (ENCODE) (Fisher exact test p-values). H) Overlaps between more accessible chromatin regions obtained by ATAC-seq after for the HCT116:PNUTS-AID cell line following the experiment indicated in Figure 1A, and HCT116 H3K9ac and H3K27ac ChIP-seq datasets (ENCODE) (Fisher exact test p-values). No significant differences were found for HCT116:Ki-67-AID and HCT116:RepoMan-AID cell lines. I) Representative western blots of histone modifications in micrococcal nuclease digested chromatin of the HCT116:Ki-67 AID (left panel), HCT116:RepoMan-AID (middle panel) and HCT116:PNUTS-AID (right panel) cell lines following the experiment indicated in Figure 1 A. The blots were probed with antibodies against H3K9me2 (first panel), H3K9ac (second panel), H3K27me3 (fourth panel), H3K27ac (fifth panel), H4K20me1 (seventh panel) and stained with total protein stain (third, sixth and eighth panel). J) Quantification of the blots in (I). The values represent the average of 3 independent biological replicas and the error bars the standard deviations. The experiments were analysed by a Student’s t-test. *= p< 0.05, **= p< 0.01, ns= not significant. K,L and M) Overlaps between less accessible chromatin regions obtained by ATAC-seq for the HCT116:Ki-67-AID (K), HCT116:RepoMan-AID (L), and HCT116:PNUTS-AID (M) cell lines following the experiment indicated in Figure 1 A, and repeat DNA sequences downloaded from UCSC genome browser using Repeatmasker. Not significant results were found in the overlaps.

To address if these changes could be a consequence of major shift in histone modifications, we analysed by immunoblotting the chromatin isolated by micrococcal nuclease digestion using several antibodies against histone modifications. Lack of Repo-Man showed a significant increase in H3K9ac and a decrease in H3K27me3 (Figure 3 I, J): these observations are congruent with our previous findings obtained using an RNAi-based analyses in HeLa cells (de Castro *et al*., 2017). Lack of Ki-67 causes a significant decrease of H3K27ac and an increase in H3K9ac while PNUTS shows only a decrease in H3K9me2 mark (Figure 3 I, J). The decrease in acetylation (H3K27ac) in Ki-67 could be related to a shift towards the more closed chromatin observed by ATAC-seq and downregulation of several genes but, for Repo-Man, the increased acetylation (H3K9ac) does not translate into a more open chromatin (at least as judged by ATAC-seq). The analyses of repeated elements within the regions that changes in the ATAC-seq experiments did not reveal any significant enrichments (Figure 3 K-M).

### PNUTS is a major regulator of transcription in G1

Among all the RIPPOs we have analysed, PNUTS behaves quite differently. First, it does not affect either the timing of mitotic exit or chromosome segregation, being the only one not to show increased aneuploidy (Figure 4 A and Supplementary Figure 3 C-F). However, we still observed a G1 block (Figure 2 I) and a moderate interferon response activation (Supplementary Figure 3 A, B). From the RNA sequencing analyses and ATAC-seq analyses, PNUTS is the PP1 targeting subunit that greatly affects the transcription landscape and chromatin accessibility (Figure 1 G, H, L and Figure 3 C). These initial findings are already very interesting because they clearly show that *in vivo*, using cell cycle-specific acute depletion, PNUTS is not involved in exiting mitosis nor in chromosome de-condesation as previous RNAi based studies had suggested. However, the data we obtained are in agreement with PNUTS involvement in transcription.

**Figure 4.**
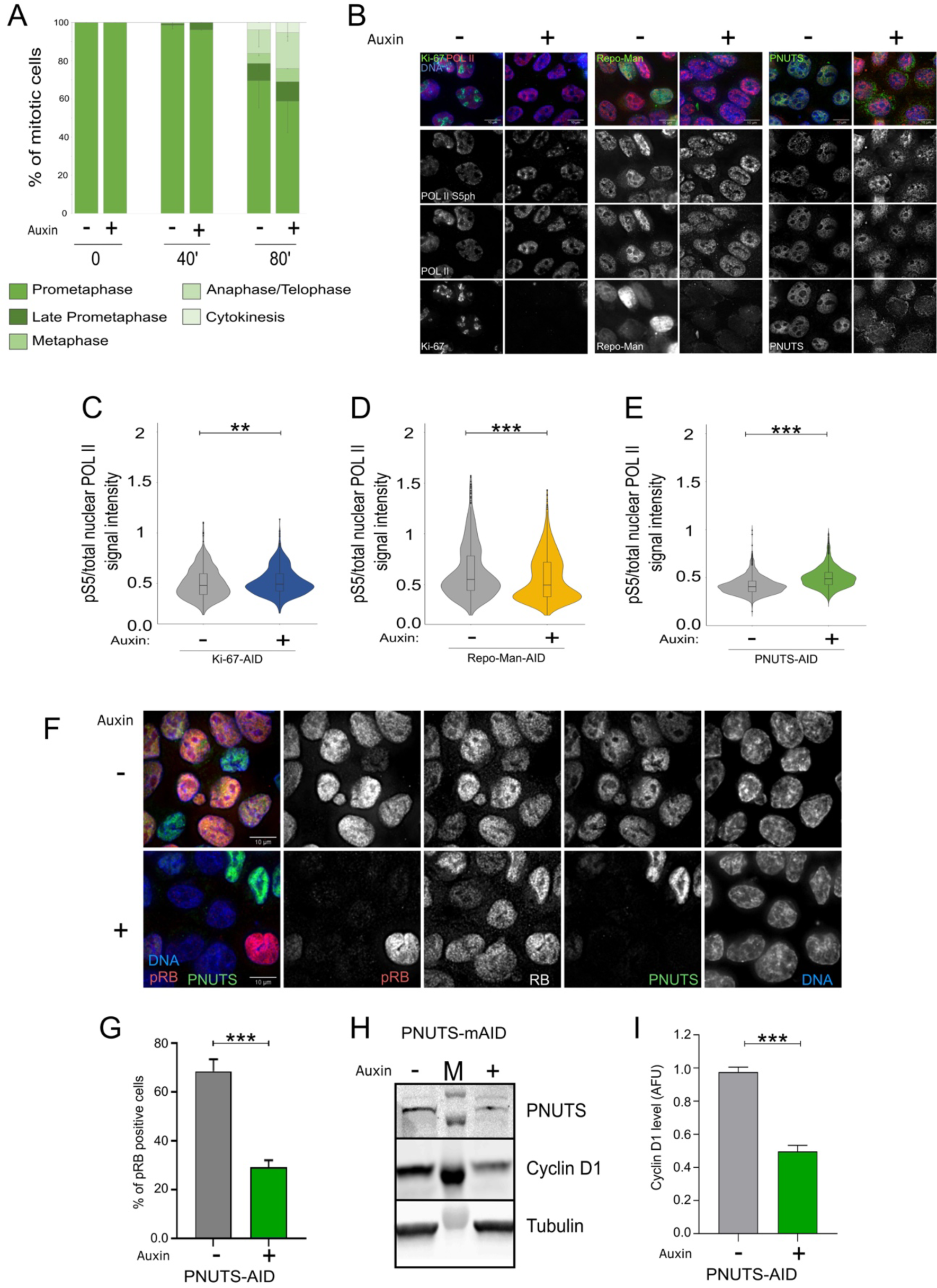
A) Quantification of different mitotic stages at 0, 40 and 80 min post-release from nocodazole in thymidine and in the presence or absence of IAA. Quantification was conducted using synchronised HCT116:PNUTS-AID cells, as described in Figure 1A, immunostained using anti-Lamin A/C and anti-Tubulin antibodies. Samples size: Control: 0=300, 40=315 and 80=322, IAA: 0=303, 40=257 and 80=354. The data were analysed by a Fisher’s exact test and the differences between Control and IAA treated cells were not significant. B) Representative images of POLII and POLII (S5ph) immunostaining using anti-POLII and anti-POLII (S5ph) antibodies on HCT116:Ki-67-AID (left panel), HCT116:RepoMan-AID (middle panel) and HCT116:PNUTS-AID cell lines treated as in Figure 1 A fixed and subjected to immunostaining, DAPI was used to stain DNA. Scale bar: 10 µm. C, D, E) Quantification of the ratio of POLII S5ph and total POLII fluorescence intensity of HCT116:Ki-67(C) AID HCT116:RepoMan-AID (D) and HCT116:PNUTS-AID (E) cell lines. The box inside the violin represents the 75th and 25th percentile, whiskers the upper and lower adjacent values and the line is the median. The data presented in the violin plots were statistically analysed with a Wilcoxon test. **=p<0.01, ***=p<0.001. F) Representative images of RB and pRB immunostaining using anti-RB and anti-pRBS807/811 antibodies on HCT116:PNUTS-AID cell line treated as in Figure 1A, fixed and subjected for immunostaining, DAPI was used as a nuclear counterstain. Scale bar: 10 µm. G) Quantification of cells positive for pRB (as in Figure 4F). The graphs represent the percentage of positive for pRB. The values are the average of 3 biological replicas and the error bars represent the standard deviations. Samples size: Control=2182, IAA=1759. The experiments were analysed by Student’s *t*-test. ***= p< 0.001 and. H) Western blot of WCL of HCT116:PNUTS-AID treated as in Figure 1A. The blots were probed for anti-PNUTs, anti-Cyclin D1 and anti-αTubulin antibodies. M: protein ladder I) Quantification of the blot in (H). The values represent the average of 3 independent replicas and the error bars are the SEM. The experiments were analysed by a Student’s t-test. ***= p<0.001,

PNUTS only established substrate is S5 in the CTD of RNAPII (pRNAPII S5) (Lee et al., 2010) (Ciurciu et al., 2013) and its dephosphorylation suppresses transcription/replication (T-R) conflicts by promoting degradation of RNAPII on chromatin, thus reducing its residence time (Landsverk *et al*., 2020). Here, we have analysed cells in G1, before replication is commenced, and we still observe a very aberrant transcription landscape with almost 2000 genes that are upregulated (Figure 1 L). We therefore wanted to test if we could detect changes in RNAPII also at this stage of the cell cycle. To this purpose we stained cells that have progressed from mitosis to G1 in the absence of PNUTS and control cells with RNAPII and RNAPIIS5ph antibodies and analysed their relative levels (Figure 4 B). These analyses show that indeed, upon PNUTS degradation, there is a significant increase of nuclear RNAPIIS5ph (Figure 4 E) supporting the idea that PNUTS is involved in the phopho-regulation of RNAPII and its absence possibly causes genome-wide acceleration of transcription that has been shown to be PP1 dependant in overexpression experiments (Cortazar et al., 2019). A decrease in this ratio is observed upon Repo-Man degradation (Figure 4 D), consistent with a halted transcription resumption.

Transcription termination defects caused by RNAPII removal are associated with R-loop formation and genome instability (Hatchi et al., 2015) (Morales et al., 2016) (Skourti-Stathaki et al., 2011). This could explain why cells that have exited mitosis without PNUTS showed an increase of damage as assessed by COMET assays, that could also be the cause for the interferon response activation (Supplementary Figure 3 G, J). Similarly, cells without Repo-Man or Ki-67 showed DNA damage at higher levels that can be explained by the presence of chromosome mis-segregation (Supplementary Figure 3 D-F and J, H, I).

Cells that have completed mitotic exit without PNUTS are also arrested in G1, although later than Repo-Man and Ki-67 depleted ones (Figure 2 E-L).

Rb protein plays a pivotal role in the negative control of cell cycle progression by blocking S-phase entry and cell growth, promoting terminal differentiation by inducing both cell cycle exit and tissue-specific gene expression (Hatakeyama and Weinberg, 1995).

This regulation is mediated by Rb association with members of the E2F1 family of transcription factors and this interaction is regulated by the phosphorylation state of Rb (Knudsen and Wang, 1997) (Calbo et al., 2002) which is controlled by the Cyclin dependent kinases (CDKs) and Protein Phosphatase 1 (PP1) (Ludlow et al., 1993).

Studies have shown that when cells are exposed to stress such as hypoxia or treatment with chemotherapeutic drugs, PNUTS dissociates from PP1, and Rb is dephosphorylated. Moreover, *in vitro* studies utilizing GST fusion constructs of PNUTS and PP1 suggest that PNUTS inhibits PP1 activity toward Rb (De Leon et al., 2008). Crystallography studies have also shown that PNUTS inhibits the PP1-mediated dephosphorylation of Rb, by blocking its binding sites on PP1 (Choy et al., 2014). To test if Rb de-phosphorylation could be the explanation for the G1 arrest of cells without PNUTS, we measured the levels of RBph (S807/ 811) in these conditions. The data clearly show that indeed Rb is significantly de-phosphorylated in cells that lack PNUTS (Figure 4 F, G). As expected, Cyclin D levels are also decreased (Figure 4 H, I).

Altogether these data suggest that PNUTS is a major regulator of transcription in G1; lack of PNUTS at this cell cycle transition leads to uncontrolled gene expression, possibly generating R loops that lead to DNA damage and prevent G1/S progression via RB dephosphorylation. However, PNUTS does not seem to play a role in mitotic exit or chromatin de-condensation as previously thought.

### Differential phosphoproteomic analyses reveal non overlapping functions for the three PP1 holocomplexes

Exiting mitosis is also accompanied by major de-phosphorylation events that are triggered both by the inhibition of mitotic kinases but also by the activation of protein Phosphatases (Vagnarelli, 2021). Repo-Man and Ki-67 are both RIPPOs that bind to PP1 stronger and more stably after anaphase onset (Booth *et al*., 2014; Kumar et al., 2016; Vagnarelli *et al*., 2011) and Repo-Man/PP1 has been shown to have a few chromosome substrates including H3T3, H3S10, H3S28 (Vagnarelli *et al*., 2011) (de Castro *et al*., 2017) (Qian *et al*., 2011) and Lamin A S22 (Huguet *et al*., 2022) while Ki-67/PP1 so far seems only to dephosphorylate itself (Takagi et al., 2014). For these reasons, we wanted to investigate the differential phosphoproteomes both in mitosis and G1 cells after the degradation of those PP1 holoenzymes. Cells were arrested in nococdazole and treated or not with IAA for 4 hrs; the two populations were SILAC labelled, and mitotic cells were fractionated into chromatin and nucleoplasmic fraction (Supplementary Figure 4 A). The samples were then processed for Mass spectrometry with and without phospho-enrichment. Other cell samples were instead released from the nocodazole block in medium containing thymidine and collected as the same time point as the previous RNA-seq and ATAC-seq experiments( Supplementary Figure 4 A)

Differentially phosphorylated proteins (cut offs –0.83 and +1.3 H/L ratios) were then compared first among the three RIPPOs. In general, this revealed little overlaps among the 3 proteins and the highest number of shared hypo- and hyper-phosphorylated proteins were between Repo-Man and PNUTS (Figure 5 A-D). The biological pathway analyses of these proteins also revealed distinct characteristics (Figure 5 E-H). Proteins that become hyperphosphorylated in M upon degradation of each RIPPO, revealed that proteins involved in gene expression processes are affected mainly by degradation of PNUTS (Figure 5 E), a finding that aligns with the transcription mis-regulation described earlier; mTOR, TRAIL signalling and interferon pathway proteins are becoming hyperphosphorylated by lack of Repo-Man and Ki-67 while the insulin pathway is unique to lack of Repo-Man (Figure 5 E). This category of proteins (Hyperphosphorylated in M) is interesting because could be enriched for mitotic substrates of these PP1 holoenzymes. The proteins that become de-phosphorylated in mitosis show more overlaps in terms of biological pathways where the 3 RIPPOs affect proteins involved in DNA replication and RNA processing (Figure 5 F). Interestingly, lack of Repo-Man seems to trigger the de-phosphorylation of proteins involved in cell cycle and the transition from Mitosis to G1 and again the insulin pathway (Figure 5 F). When the analysis is conducted for proteins that become differentially phosphorylated in G1, PNUTS seems the main RIPPO that, when missing during mitotic exit, leads to a significant enrichment in a few categories, including gene expression and CDC42 regulation (Figure 5 G). For the hypophosphorylated category, the RIPPO affecting several categories is Ki-67 while PNUTS affects the phosphoregulation of transcription processes, including C-MYC targets and Repo-Man the insulin signalling (Figure 5 H). These findings suggest a dominant role of PNUTS in transcription regulation at this cell cycle stage (which aligns with some of the reported findings of PNUTS - (Landsverk *et al*., 2020) and highlight a previously unknown involvement of Repo-Man in the Insulin pathway regulation and glucose signalling.

**Figure 5.**
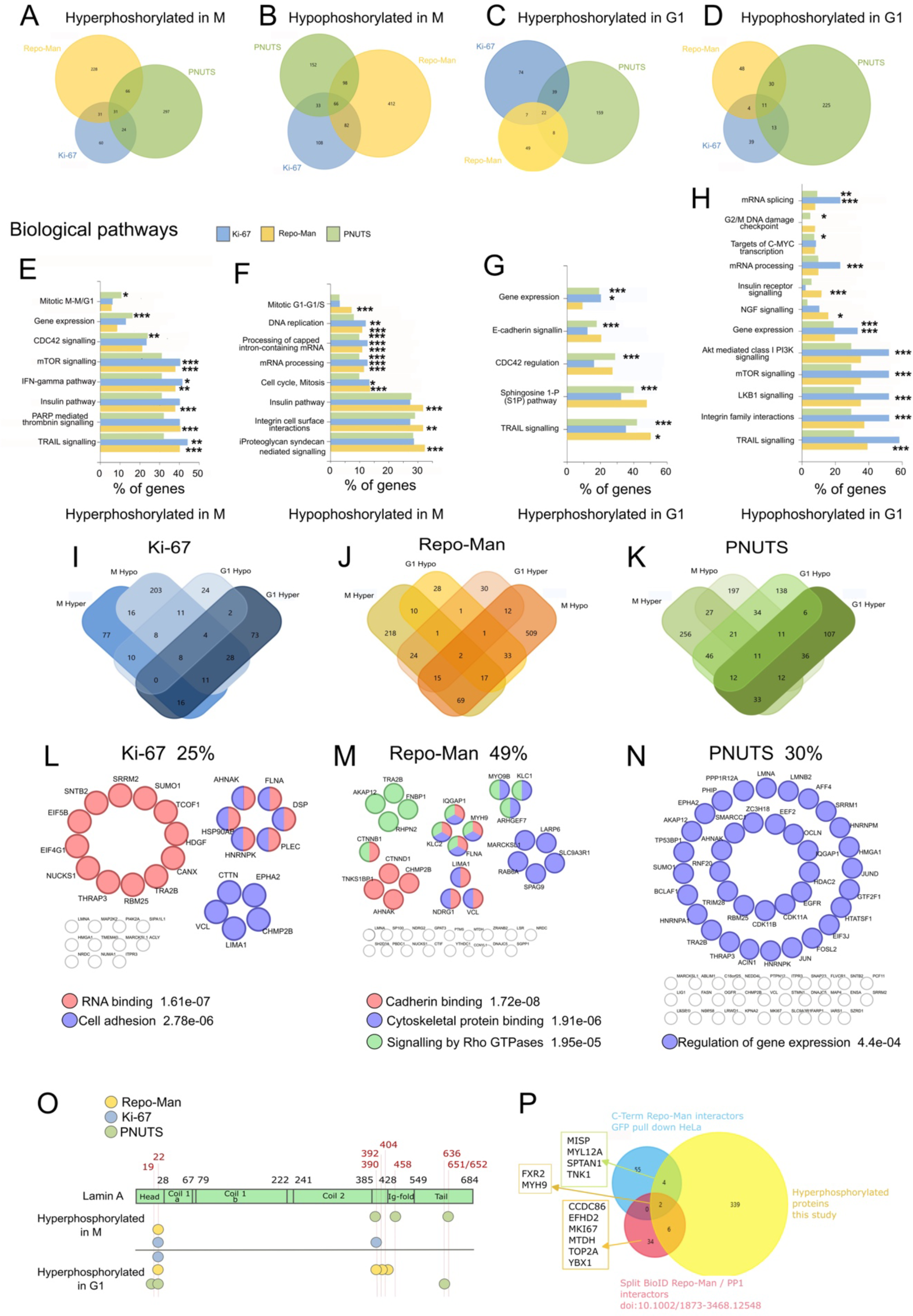
A and B) Venn diagrams of SILAC-based phospho-proteomic analyses in Mitosis of Hyperphosphorylated proteins (A) and Hypophosphorylated proteins (B) for HCT116:Ki-67-AID, HCT116:RepoMan-AID and HCT116:PNUTS-AID cell lines treated with or without IAA according to the scheme in Supplementary Figure 3 A. C and D) Venn diagrams of SILAC-based phospho-proteomic analyses G1 for Hyperphosphorylated proteins (C) and Hypophosphorylated proteins (D). Samples collected for phosphoproteomic analyses from HCT116:Ki-67-AID, HCT116:RepoMan-AID and HCT116:PNUTS-AID cell lines treated with or without IAA according to the scheme in Supplementary Figure 3 B. E, F, G and H) FunRich biological pathways enrichment analysis of the Hyperphosphorylated (E) and Hypophosphorylated (F) proteins in Mitosis and Hyperphosphorylated (G) and Hypophosphorylated (H) proteins in G1 I, J and K) Venn diagram of HCT116:Ki-67-AID (I), HCT116:RepoMan-AID (J) and HCT116:PNUTS-AID (K) Hyperphosphorylated and Hypophosphorylated proteins in Mitosis and in G. L,M and N) String analyses of proteins that remain hyperphosphorylated from Mitosis to G1 for HCT116:Ki-67-AID (L), HCT116:RepoMan-AID (M) and HCT116:PNUTS-AID (N) O) Scheme of Lamin A protein structure and domains with Hyperphosphorylated or Hypophosphorylated residues found in the phosphoproteomics analyses of HCT116:Ki-67-AID (blue), HCT116:RepoMan-AID (yellow) and HCT116:PNUTS-AID (green) cell lines in Mitosis (M) and G1. P) Venn diagram of hyperphosphorylated proteins of HCT116:RepoMan-AID cell line in Mitosis, GFP-RepomanC-term interactors GFP pull-down Hela and Split BioID Repo-Man / PP1interactors (De Munter *et al*., 2017).

The category of hyperphosphorylated proteins in G1 (Figure 5 G) should contain substrates of these holoenzymes during mitotic exit; however, the hyperphosphorylation could also be the aftermath of mitotic substrates that become hyperphosphorylated before exiting mitosis and then maintaining a higher phosphorylation level throughout G1. To understand how many proteins belong to this category, we analysed the overlaps for each RIPPO (to be noted that we did not focus on a specific phosphosite at this stage). Interestingly, between 25 to 49 % of hyperphosphorylated proteins in G1 are also hyperphosphorylated in M (Figure 5 I-N). These findings are important and need to be taken into consideration when analysing phosphoproteomic data that are collected either by RNAi approaches or by protein degradations and when the depletion of a protein could affect other cell cycle stages. Among these proteins, again PNUTS shows a significant enrichment for proteins involved in transcription and parallels nicely the major transcription dysregulation observed in G1 (Figure 5 N). For Ki-67, cell adhesion proteins are part of the category (Figure 5 L); this is also interesting in the light of the upregulation of MET genes (Supplementary Figure 3 J) and the results obtained by the Fisher’s lab in 4T1 cells showing that loss of Ki-67 deregulates Pathways Involved in Cancer (Mrouj *et al*., 2021).

These datasets provide an excellent resource, as they are enriched of possible substrates, but of course, they also contain proteins whose phosphorylation could differ as secondary event. However, intersecting RIPPOs pull down experiments or proximity biotinylation approaches with these differential phosphoproteomes, would provide a list of highly probable substrates (Kommer et al., 2024). We have conducted the following analyses with Repo-Man because we have previously performed pull-down experiments in human cells (HeLa) and the Bollen lab has conducted biotinylation experiments using a Split BioID approach in HEK293T cell (De Munter et al., 2017). The intersects of these datasets have revealed some interesting possible substrates for Repo-Man/PP1 including Myosin Heavy Chain 9 (MYH9 – with a role in cytokinesis), Topoisomerase IIα, Ki-67 and CCDC86 (involved in chromosome structure and segregation) (Figure 5 P). The identification of Ki-67 as possible Repo-Man substrate is also quite interesting and could explain why some of the changes in G1 observed upon Repo-Man degradation overlap with the ones found upon Ki-67 degradation.

The analyses described above, mainly focussed on proteins unique for each RIPPOs as you could assume that one holoenzyme is specific for the de-phosphorylation of a single protein. However, some common proteins could be phosphorylated in multiple sites and each site could also be de-phosphorylated at different stages by different proteins. As an example, we have analysed the phosphorylation status of Lamin A because it appears in all the samples, and it is also a Repo-Man substrate (Huguet *et al*., 2022) (Moriuchi and Hirose, 2021). Several Lamin A sites become hyperphosphorylated in mitosis upon degradation of each RIPPO: S22 (Ki-67 and Repo-Man), S390 (PNUTS and Ki-67), S494 (PNUTS), and 651 (PNUTS). In the G1 phosphoproteome, the S19 and S636 are only found when PNUTS is degraded, while 390, 392 and 404 are only found upon Repo-Man degradation and these sites are important for the regulation of the nuclear localisation of Lamin A and its polymerisation (Kochin et al., 2014) (Makarov et al., 2019) (Figure 5 O); these findings indicates that Repo-Man is the major Lamin A phosphatase.

In summary, these comparative analyses show no redundancy for the three complexes as they are characterised by quite distinct patters of phosphorylation changes: hyperphosphorylations in mitosis was more prominent upon degradation of Repo-Man and PNUTS but not of Ki-67, whereas degradation of Repo-Man caused major changes in the de-phosphorylation of several protein (which are clearly downstream effects of some important biological functions of Repo-Man in mitosis).

### Ki-67 is essential at the M/G1 transition to maintain centromere organisation

The discovery that lack of Ki-67 during mitotic exit leads to aneuploidy, a block at the restriction checkpoint together with previous reports showing that Ki-67 promotes timely replication of centromeres (van Schaik *et al*., 2022) and centromeric organisation in S phase (Stamatiou *et al*., 2024) led us to investigate if a link between centromeres organisation and Ki-67 could be important at this cell cycle stage. Using the differential phosphoproteome datasets for Ki-67 in mitosis, we conducted STRING analyses and revealed a significant enrichment for proteins that are important for chromosome organisation, cell division but also centromeric proteins (Figure 6 A, Supplementary Figure 4 C); among these, we identified changes in phosphorylation for Sororin (CDCA5), which is essential for the maintenance of chromatid cohesion in mitosis (Ladurner et al., 2016); Survivin (CDCA3), necessary for the recruitment of the CPC to centromeres (Jeyaprakash et al., 2011); NUF2, a component of the NDC80 complex and CENP-C, a key kinetochore components which phosphorylation promotes kinetochore assembly (Hinshaw et al., 2023). We therefore wanted to assess the centromeric organisation of mitotic chromosomes depleted of Ki-67. For this, cells were arrested in Nocodazole and Ki-67 was degraded once cells have already entered mitosis. We first conducted FISH for the centromeric satellites of chromosomes 1 and then measured the area occupied by the probes to test if changes of heterochromatin compaction could be detected at this stage (we did see it happening upon replication without Ki-67 (Stamatiou *et al*., 2024). The analyses show that, when Ki-67 is degraded in mitosis, these centromeric regions are expanded (Figure 6 B), suggesting that Ki-67 could play a role in their organisation. We also conducted the same experiments in cells that have exited mitosis without Ki-67 and obtained the same results for both chromosome 1 and 15 (Figure 6 C). To be noted that the total area of the nucleus is not becoming bigger in the same conditions, but it is significantly smaller (Figure 6 D), thus ruling out a more general role of Ki-67 on chromatin compaction. All these parameters were not affected by degradation of Repo-Man or PNUTS (Supplementary Figure 5 A-D), thus suggesting a specific role for Ki-67 at the pericentromeric chromatin. Previous work has shown that centromere decompaction and increased in the total area of centromeric DNA is associate with lack of CENP-B (Chardon et al., 2022). We therefore analysed the level of the centromeric protein CENP-B in these cells and we could show a decrease of CENP-B at the centromeric regions both in mitosis (Figure 6 E) and in G1 (Figure 6 F) (however its transcription is not decreased (Supplementary Figure 5 E)). Altogether these data indicate that, without Ki-67, the organisation of centromeric chromatin is compromised and leads to a decrease of CENP-B. Ki-67 has never been reported to be at the centromere in mitosis but it accumulates at the periphery of the chromosomes (albeit in a dynamic manner (Saiwaki et al., 2005)); we therefore wanted to evaluate at higher resolution if a subpopulation of Ki-67 was indeed visible at the centromeres in mitosis and G1. We prepared chromosome spreads and stained for Ki-67 and ACA: here we could detect a partial overlap between Ki-67 and the centromeric proteins (Figure 6 G); to confirm this, we used a proximity ligation assay (PLA) to assess further the close localisation between Ki-67 and CENP-B; also in this case we found positive PLA signals in mitosis only when Ki-67 was present (Figure 6 I). We then analysed the localisation of endogenous Ki-67 and CENP-B in early G1; in these cells that have not yet reformed the nucleoli, Ki-67 is present in aggregates and CENP-B is in proximity of these localised Ki-67 clusters where Ki-67 seems to enwrap the centromeres (Figure 6 H) and, when analysed by DeepSIM, we could clearly see co-localisation between Ki-67 and CENP-B in these cells. Analyses of mitotic exit showed that cells lacking Ki-67, slightly delay in chromosome alignment (Supplementary Figure 5 F): this in in agreement with spindle assembly defects previously observed in Ki-67 knock-down cells (Cuylen et al., 2016) but, eventually, all the cells exit mitosis (Figure 1 B, C, D – G1) although with chromosome segregation errors (Supplementary Figure 5 G). Therefore, these analyses led us to conclude that Ki-67 is in proximity of the centromeres from mitosis to early G1 and it is important for maintaining both a compact centromere structure, a normal phosphorylation of a subset of centromeric proteins, and CENP-B levels. These novel findings showing a link between Ki-67 and centromeres can also explain why degradation of Ki-67 in mitosis leads to aneuploidy (Supplementary Figure 3 D-F).

**Figure 6.**
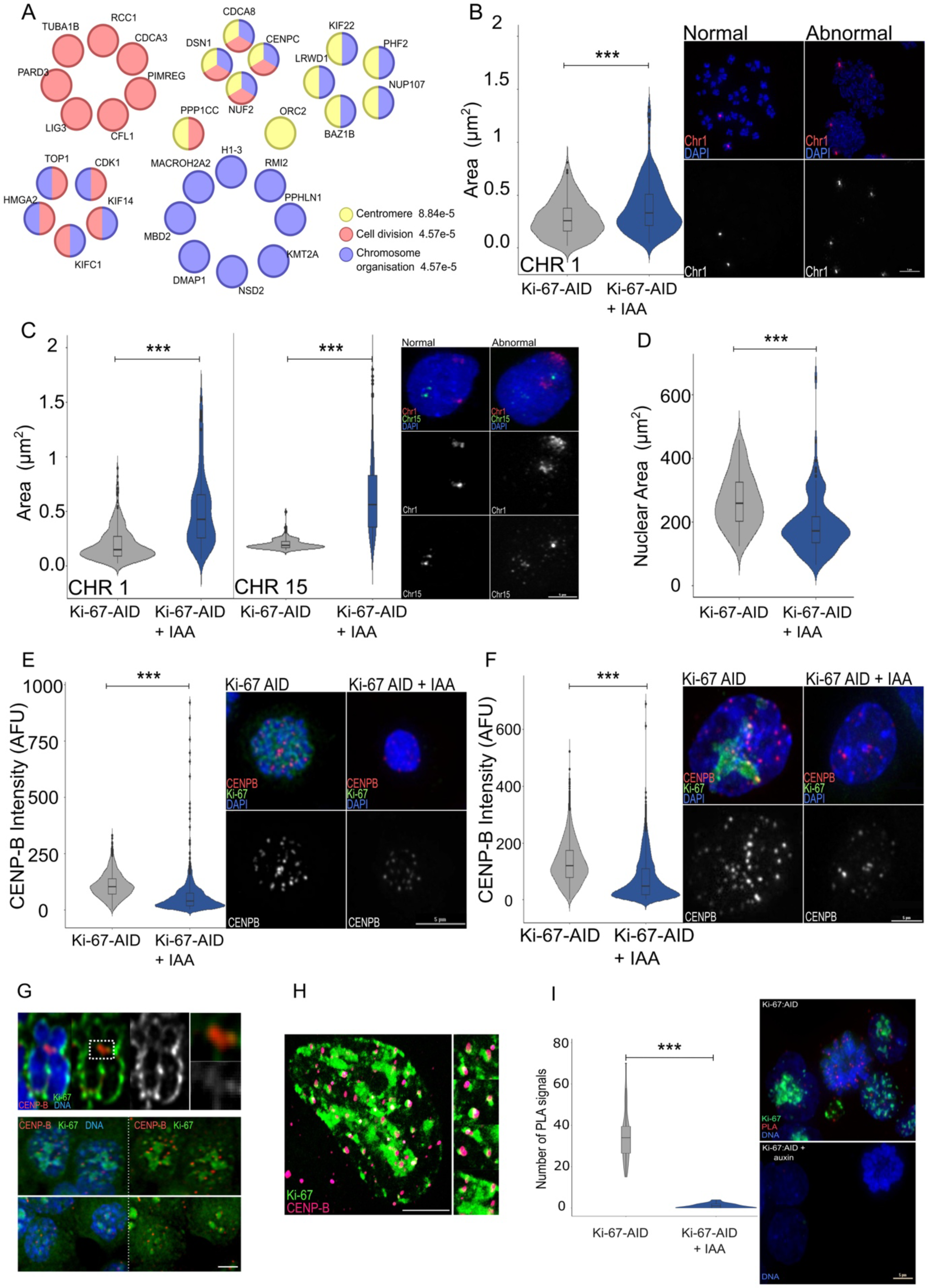
A) String analysis of differentially phosphorylated proteins from Figure 5 B for HCT116:Ki-67-AID cell line. B) HCT116:Ki-67-AID were treated with or without IAA for 4 h after 18 h incubation with nocodazole, fixed and subjected to FISH with a centromeric chromosome 1 probe. Representative images of the FISH normal and abnormal signals and violin plot representing the quantification of the area occupied by the FISH signals (left). Scale bar 5 μm. Sample size: Control=194, IAA=296. The box inside the violin represents the 75th and 25th percentile, whiskers the upper and lower adjacent values and the line is the median. The data presented in the violin plots were statistically analysed with a Wilcoxon test. ***=p<0.001 C) HCT116:Ki-67-AID cells were treated as indicated in Figure 1A, fixed and subjected to FISH. Representative images of the FISH normal and abnormal signals obtained with the pUC 177 (red signals, Chr 1) and pTRA-20 (green signals, Chr 15) probes (right) and quantification of the area occupied by the FISH signals (middle and left). Scale bar 5 μm. Sample size: CHR1: Control=280, IAA=284 and CHR15: Control=253, IAA=235. The box inside the violin represents the 75th and 25th percentile, whiskers the upper and lower adjacent values and the line is the median. All the data presented in the violin plots were statistically analysed with a Wilcoxon test. ***=p<0.001 D) HCT116:Ki-67-AID were treated as in Figure 1 A and fixed. Quantification of the nuclear area in μm2. Scale bar 5 μm. Sample size: Control=95, IAA=114. The box inside the violin represents the 75th and 25th percentile, whiskers indicate the upper and lower adjacent values and the line is the median. The data presented in the violin plots were statistically analysed with a Wilcoxon test. ***=p<0.001 E) Representative images of CENP-B immunostaining using anti-CENP-B antibody on HCT116:Ki-67-AID treated with or without IAA for 4 h after 18 h incubation with nocodazole (right) and quantification of CENP-B intensity (left). Scale bar 5 μm. Sample size: Control=2139, IAA=1646. The box inside the violin represents the 75th and 25th percentile, whiskers the upper and lower adjacent values and the line is the median. The data presented in the violin plots were statistically analysed with a Wilcoxon test. ***=p<0.001 F) Representative images of CENP-B immunostaining using anti-CENP-B antibody on HCT116:Ki-67-AID were treated as in Figure 1 A (right) and quantification of CENP-B intensity (left). Scale bar 5 μm. Sample size: Control=1786, IAA=3985 foci. The box inside the violin represents the 75th and 25th percentile, whiskers the upper and lower adjacent values and the line is the median. The data presented in the violin plots were statistically analysed with a Wilcoxon test. ***=p<0.001 G) Representative images of CENP-B immunostaining using anti-CENP-B antibody on HCT116:Ki-67-AID. Mitotic chromosomes (top), early G1 (bottom). Scale bar 5 μm. H) Representative images of CENP-B immunostaining using anti-CENP-B antibody on HCT116:Ki-67-AID. Early G1 by Deep SIM microscopy (right). Scale bar 10 μm I) Representative images Proximity ligation assay (PLA) using anti-CENP-B and anti-GFP antibodies on asynchronous HCT116:Ki-67 AID cell line without (top) of with (bottom) IAA (right) and quantification of PLA signals in mitotic cells (left). Scale bar 5 μm. The box inside the violin represents the 75th and 25th percentile, whiskers the upper and lower adjacent values and the line is the median. Sample size: Control=36, IAA=22. The data presented in the violin plot were statistically analysed with a Wilcoxon test. ***=p<0.001.

These analyses altogether suggest that, although both Ki-67 and Repo-Man degradation leads to aneuploidy, an interferon response activation and G1 arrest, a centromeric defect is only Ki-67 specific, placing Ki-67 in a unique signalling pathway. It would be interesting to investigate if Ki-67 itself is an important effector of the signalling cascade that links aneuploidy and centromere structure defects to the interferon response activation and if it could be a signalling trigger for cell cycle exit (quiescence) or senescence.

### Repo-Man is essential for maintaining the checkpoint in mitosis

Our analyses have also shown that a mitotic exit without Repo-Man caused an increase in aneuploidy, G1 arrest and interferon signalling activation but it is not associated with centromeric chromatin defects. We therefore wanted to analyse in more detail the phosphoproteome of mitotic cells where Repo-Man has been degraded.

We extracted the differentially Repo-Man specific phosphoproteins and conducted STRING analyses. Without Repo-Man, several cell cycle and cell-cycle mitotic proteins presented differential phosphorylation when Repo-Man was degraded in mitosis together with the de-phosphorylation of a significant proportion of APC/C components, including CDC20 S41 (Figure 7 A, Supplementary Figure 4 D). Clearly, this scenario suggests that Repo-Man could be important for maintaining cells in mitosis, a role that so far has not been defined for this protein and it is against the current assumption that actually Repo-Man is inactivated in mitosis and mainly needed to direct mitotic exit specific dephosphorylation events (Vagnarelli *et al*., 2011) (Qian et al., 2013).

**Figure 7.**
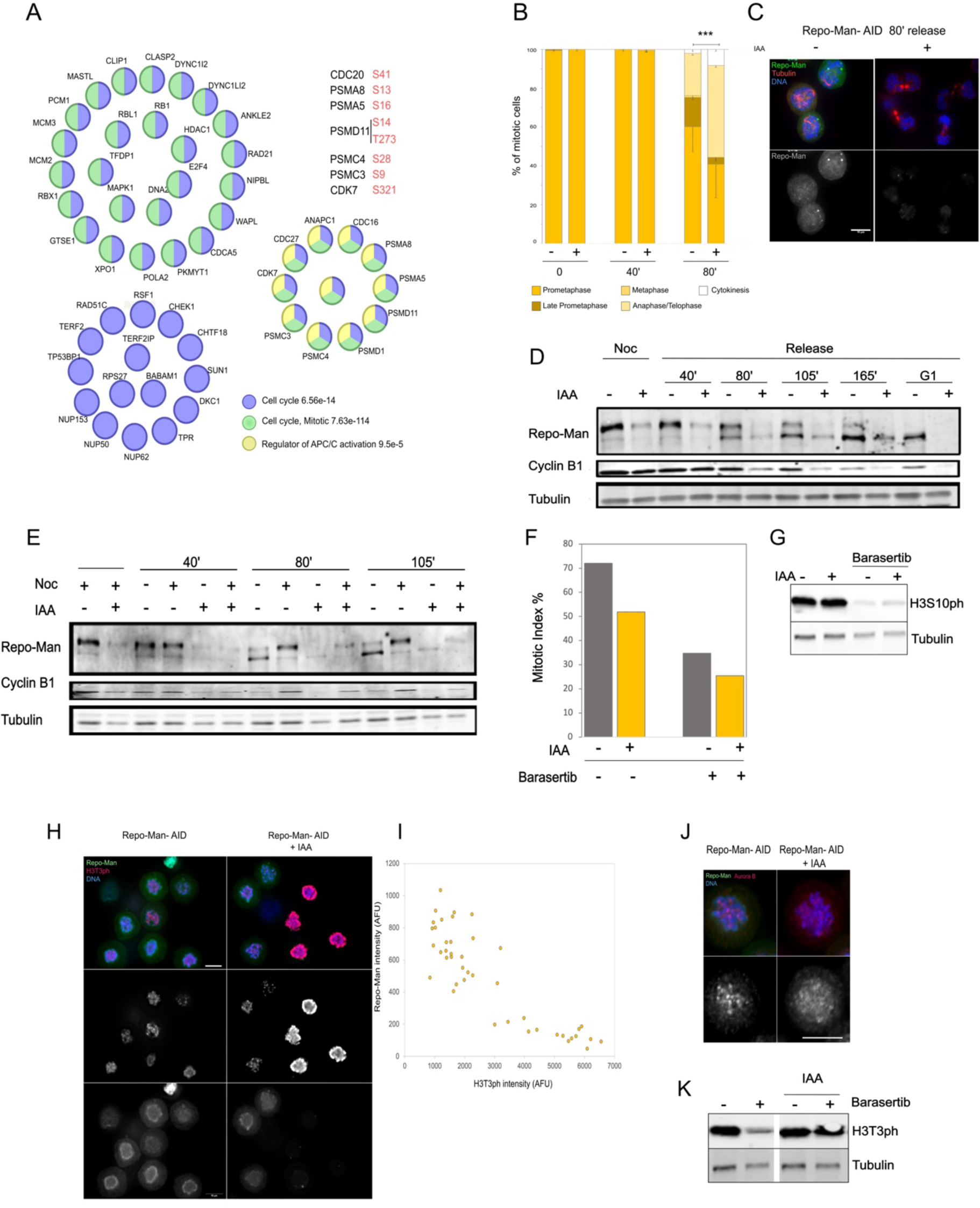
A) String analysis of differentially phosphorylated proteins from Figure 5 B for HCT116:RepoMan-AID cell line. B) Quantification of different mitotic stages at 0, 40 and 80 min post-release from nocodazole in thymidine and in the presence or absence of IAA. Quantification was conducted using synchronised HCT116:RepoMan-AID cells, as described in Figure 1A, immunostained using anti-Lamin A/C and anti-Tubulin antibodies. Samples size: Control: 0=194, 40=208 and 80=199, IAA: 0=200, 40=203 and 80=165. The data were analysed by a Fisher’s exact test, ***=p<0.001. C) Representative fluorescent microscopy image of HCT116:RepoMan-AID cells 80 min post release from nocodazole, in the absence or presence of IAA, immunostained using anti-αTubulin antibody. Scale bar 10 μm. D) HCT116:RepoMan-AID treated as in Figure 1A. Cells were collected 40, 80, 105, 165 min or overnight after the release from nocodazole, in the absence or presence of IAA. Whole cell lysates were blotted with anti-Repo-Man (top panel), anti-Cyclin B1 (middle panel) and anti-αTubulin (bottom panel) antibodies. E) HCT116:RepoMan-AID cells treated as in Figure 1A or kept in nocodazole. Cells were collected 40, 80 or 105 min after the release from nocodazole, in the absence or presence of IAA. Whole cell lysates were blotted with anti-Repo-Man (top panel), anti-Cyclin B1 (middle panel) and anti-αTubulin (bottom panel) antibodies. F) Quantification of the mitotic index of HCT116:RepoMan-AID cells treated with nocodazole 18 h, before the addition of IAA for 4 h in the absence or presence of Barasertib. G) Western blot of WCL of HCT116:RepoMan-AID cells treated with nocodazole 18 h, before the addition of IAA for 4 h in the absence or presence of Barasertib. The blots were probed with anti-H3S10ph (top panel) and anti-αTubulin (bottom panel) antibodies. H) Representative images of H3T3ph immunostaining using anti-H3T3ph antibody on HCT116:RepoMan-AID cells treated with nocodazole 18 h, before the addition of IAA for 4 h. I) Scatter plot representing the intensity of Repo-Man and H3T3ph from the experiment in (H) R^2^=0.756. J) Representative images of Aurora B immunostaining using anti-Aurora B antibody on HCT116:RepoMan-AID cells treated with nocodazole 18 h, before the addition of IAA for 4 h. scale bar= 10 μn. K) Western blot of WCL of HCT116:RepoMan-AID cells treated with nocodazole 18 h, before the addition of IAA for 4 h in the absence or presence of Barasertib. The blots were probed with anti-H3T3ph (top panel) and anti-αTubulin (bottom panel) antibodies.

Taking advantage of the degron system we generated, we could test this hypothesis. First, we repeated the mitotic exit experiments where cells were blocked in nocodazole, Repo-Man is degraded and then cells are released in medium with and without IAA. Mitotic exit progression was monitored by tubulin staining. As the distribution of mitotic stages indicate, at 80 min, 70% of the cells with Repo-Man are still in prometaphase/metaphase while 50% of the cells without Repo-Man have progressed already to anaphase/telophase (Figure 7 B, C). To validate this, we analysed the degradation of Cyclin B in the same time course, and we obtained similar results with Cyclin B being degraded at a faster rate in cells without Repo-Man (Figure 7 D). We then asked if this effect was also occurring in cells blocked in nocodazole, where the checkpoint is very strong. To test this, we blocked the cells in nocodazole overnight, degraded Repo-Man and monitored both the Cyclin B levels but also the mitotic index. Even in this case, the level of Cyclin B decreases in the absence of Repo-Man (Figure 7 E) and so the mitotic index (Figure 7 F, left bars). These data clearly indicate that Repo-Man is necessary for maintaining cells in mitosis. We and others have previously shown that depletion of Repo-Man by RNAi leads to a mislocalisation of H3T3ph in mitotic cells and suggested that this phosphosite is a substrate of the complex. However, these experiments could not discriminate if the mislocalisation was caused by a function of the complex at mitotic entry or it indicated a requirement for Repo-Man throughout mitosis. We therefore tested this by degrading Repo-Man when cells are already in mitosis and evaluate the level and distribution of H3T3ph. As the data show, degradation of Repo-Man in mitosis causes a major increase in H3T3ph levels but also its spreading along the chromosome arms (Figure 7 H-I); this is in agreement with previous results but also clearly shows that Repo-Man/PP1 complex is active in mitosis. Interestingly, in cells where Repo-Man has not been effectively degraded, we can see a clear correlation between Repo-Man levels and H3T3ph levels (Figure 7 I). Moreover, as expected, Aurora B localisation was impaired (Figure 7 J). We then wondered if the role of Repo-Man in mitosis is purely to maintain Aurora B activity. We tested this by checking the mitotic index for cells arrested in Nocodazole followed by degradation of Repo-Man and inhibition of Aurora B by the highly selective inhibitor Barasetib (Johnson et al., 2023): an experiment impossible to conduct without the Repo-Man degron cell line we generated; here the hypothesis was that if Repo-Man is working on the same pathway of Aurora B, inhibition of Aurora B in the presence or absence of Repo-Man would have the same effect. However, this was not the case because degradation of Repo-Man has an additive effect on Aurora B inhibition (Figure 7 F), thus suggesting that Repo-Man might also affect another pathway. The efficient inhibition of Aurora B was also monitored by analysing the level of its known mitotic substrate H3S10ph (an Aurora B substrate) by western blotting (Figure 7 G). We then finally checked the level of H3T3ph upon inhibition of Aurora B in the presence or absence of Repo-Man; here we could see a decrease of H3T3ph upon inhibition of Aurora B as previously reported: this had been interpreted as the results of inhibition of Haspin kinase that requires Aurora B for its activity (Wang et al., 2011). Surprisingly, we did see a rescue of the phosphorylation levels when we degrade Repo-Man and inhibit Aurora B (Figure 7 K). These results are not consistent with the current model that includes Haspin activity being dependant on Aurora B. The results obtained instead indicate that Haspin activity does not depend on Aurora B, and that the effect of Aurora B inhibition on H3T3 levels (decreased) is mediated by the re-localisation of Repo-Man to the chromatin (Qian *et al*., 2013).

Altogether this unbiased approach has revealed an essential role for Repo-Man in maintaining the mitotic checkpoint. Future work will unveil how this Repo-Man function intersects with the SAC regulation.

## Discussion

The identification of specific roles and contributions of proteins in cell cycle progression have been hindered by the intrinsic limitation of methodologies that were not targeted enough to only affect a particular cell cycle transition. Although these approaches have allowed the discovery of many important features of cell cycle, the relationship between the effects described by the depletion of a protein and the independence from the roles they elicited in other cell cycle stages has not been possible. The availability of protein-degradation tags that can lead to a rapid depletion of a protein when cells are in a specific cell cycle stage, has allowed to revisit some of these functions. In addition, lots of the discoveries have been hypothesis-driven and based on previous knowledge; this has been extremely important in identifying detailed molecular mechanisms, but they have limited the exploration and discovery of unknown and unpredicted functions.

Here, we have combined a protein rapid degradation approach with unbiassed multi-omics to investigate the role of 3 major chromatin-associated PP1 targeting subunits in mitotic exit progression and G1 nuclear programme establishment.

We have endogenously tagged with AID degron modules Repo-Man, PNUTS and used the previously described Ki-67-AID cell line (Stamatiou *et al*., 2024). The proteins have been degraded when cells were already in prometaphase (thus eliminating all the problems that could be related to S phase, DNA damage repair and mitotic entry) and then allowed to progress through mitotic exit. We then followed 3 major events that are linked to mitotic exit: 1) transcription resumption using RNA seq; 2) chromatin re-opening by assessing chromatin accessibility using ATAC-seq and 3) protein de-phosphorylation using differential phosphoproteomic. This represents an unbiassed approach that, besides providing the community with important datasets, led us to unveil some specific functions of these RIPPOs at this cell cycle transition.

This approach first revealed that there is no redundancy in the system and each protein has a very specific profile even if some sequence similarities and phylogeny could have hypothesized that Ki-67 and Repo-Man could have complemented each other (Booth *et al*., 2014).

We have also shown that all these RIPPOs are essential for a faithful mitotic exit and G1 re-establishment, and lack of each one leads to a G1 block as shown by the transcriptomic profiles in G1 cells and the analyses of CDK2 and APC/C^CDH1^ reporters. This is the first time that such an effect has been shown and clearly separated by other cell cycle problems. The mitotic exit time course allowed us to reveal that for Ki-67 and Repo-Man, the major defect driving the block is linked to chromosome mis-segregation: high levels of aneuploidy were observed upon a mitotic exit in the absence of these proteins. This is again an aspect that has never been identified previously. However, the missegregation defects are linked to different molecular mechanisms; we have shown that for Repo-Man they are caused by a premature mitotic exit and attenuation of the SAC but, for Ki-67, defects in the centromere chromatin structure are the leading cause. Interestingly, using this method, we could discard a prominent role of PNUTS in either maintaining or exiting mitosis as was previously suggested (Wang et al., 2019). Moreover, we did not observe any changes in chromatin de-condensation in this *in vivo* system, making it unlikely that PNUTS plays a role in chromatin de-condensation (Landsverk *et al*., 2005), actually we see a more accessible chromatin as shown by the ATAC seq analyses.

Changes in the transcriptomic profiles for Repo-Man and Ki-67 are mainly linked to the G1 checkpoint arrest rather than to changes in chromatin accessibilities, in fact the ATAC seq analyses did not reveal major changes of this parameter.

On the other hand, we could show that PNUTS plays a major role in regulating transcription at this cell cycle transition where its role is to refrain uncontrolled transcription. Molecularly, we could identify increased nuclear levels of RNAPII S5ph that have been reported previously upon PNUTS knockdown and during replication; here we show that this imbalance already occurs in G1. We also observed hyperphosphorylation of Rb in cells that exited mitosis without PNUTS which could explain the late G1 block seen in these cells.

Mitotic exit without each of these proteins also triggered an interferon response; this was expected for Repo-Man and Ki-67 due to aneuploidy but PNUTS lacking cells are not aneuploid. This scenario gave us the opportunity of comparing the two responses and possibly identifying and distinguish an interferon signature for the aneuploidy-induced response and a transcription mis-regulation-linked one. Genes such IFITM1, OAS2, PSMB8, MX1, ITGB3 and IRF7 are common to both system but IFIT1B, IFIT3, RIOK3, CFB, DHX58, HLA-C, CDC274, CLU, LILRB1, PSMB9 and GAB2 are only shared between Repo-Man and Ki-67. It would be interesting in the future to evaluate these gene regulations in situations of chronic aneuploidy such as Down syndrome but also to understand their role in CIN driven cancers.

The differential phosphoproteomic analyses gave us also some important clues on how such datasets should be interpreted. First, we observed several differentially hyperphosphorylated proteins when cells exited mitosis in the absence of each protein. However, a closer comparison with the differential prometaphase phosphoproteins revealed that a high percentage of these proteins were common. This data show that caution should be exerted when analysing these types of datasets; in fact, although they could reveal interesting substrates, they do not really represent bona fide mitotic exit targets but just a legacy from the mitotic ones. Omitting this distinction could lead to very different interpretations and should be taken into account in future investigations.

The phosphoproteomic analyses in mitosis also provided us with more and unexpected clues on the mitotic functions of Repo-Man and Ki-67.

Repo-Man has been shown to be important for de-phosphorylating H3T3, H3S28, H3S10 in mitotic exit and its activity was thought to be negatively regulated by Aurora B and CDK1 in early mitosis. Here we show that Repo-Man is essential for maintaining the SAC signalling; its degradation in fact leads to a premature mitotic exit as evaluated by Cyclin B degradation and decreased mitotic index. This effect seems not to be solely mediated by Aurora B function and more investigations will be able to understand mechanistically how this is achieved. Moreover, we have unveiled new APC/C phosphosites that are linked to its activation that were so far not known. These data therefore set new parameters: 1) Repo-Man is very active in M phase, 2) it is required for the SAC and 3) Haspin kinase (the kinase Repo-man counteracts in mitosis) is not regulated by Aurora B. New models are therefore required to explain how the centromeric H3T3ph is protected from a very mitotically active Repo-Man.

The other discovery this unbiassed approach allowed us to reach is related to the link between Ki-67 and centromeric maintenance. The differential phosphoproteomic has revealed that several centromeric proteins, including CENP-C change phosphorylation status upon Ki-67 degradation in prometaphase. Phosphorylation of CENP-C by Aurora B has been shown to facilitate kinetochore attachment error correction in mitosis (Zhou et al., 2017) but also CDK1-mediated CENP-C phosphorylation seems to modulate CENP-A binding and mitotic kinetochore localization (Watanabe et al., 2019): these changes per se could explain the aneuploidy we observed upon Ki-67 degradation. We and others have reported defects in centromeric chromatin replication and structure upon Ki-67 degradation during S phase (van Schaik *et al*., 2022) (Stamatiou *et al*., 2024) and older observations have reported Ki-67 co-localisation with satellite DNA in G1 (Bridger et al., 1998). Here we show for the first time that Ki-67 is indeed in close proximity of the centromere in mitosis and its degradation affects centromeric compaction and CENP-B binding. The localisation of Ki-67 close to centromeric chromatin is still maintained in early G1 as we show by super resolution. These new findings are very interesting on different aspects. First, RNAi depletion of chromosome periphery proteins that are recruited by Ki-67 have been linked to chromosome segregation defects (Fujimura et al., 2020) (Amin et al., 2007; Ma et al., 2007; Stamatiou et al., 2023) (Ugrinova et al., 2007) but Ki-67 null cells, apparently, did not present segregation defects (Cuylen *et al*., 2016); this discrepancy could be due to either adaptation mechanisms of not completed depletion using RNAi methods. Here we show that cells acutely depleted of Ki-67 do show mis-segregation defects.

Interestingly, Ki-67 has also been suggested to be the homologue in metazoan of “bridging”, a kinetochore component identified in Cryptococcus neoformans that connects outer kinetochore to the centromeric chromatin (Sridhar et al., 2021), but so far no reports have ever associated Ki-67 with kinetochores. Our new findings show that this seems to be the case. It would be therefore very interesting to verify how these two related proteins are working to maintain a functional kinetochore across species.

## Acknowledgment

The authors thank Prof A.Sala (Brunel) and E.Schirmer (Edinburgh) for critical discussions and suggestions.

The work was supported by the Wellcome Trust Investigator Award 210742/Z/18/Z to Paola Vagnarelli; KS was supported by a CHMLS PhD scholarship (Brunel University London).

The Proteomic work was conducted at the Wellcome Discovery Research Platform for Hidden Cell Biology [Wellcome grant number 226791].

## Author Contribution

Conceptualisation: PV, CS; Methodology: KS, FH, MB, CS, JR, PV; Formal Analysis: KS, FH, MB, IJdC, CS, PV; Inves;ga;on: KS, FH, MB, IJdC, CS, PV; Data Cura;on: KS, FH, MB, CS, JR, PV; Wri;ng – Review & Edi;ng: PV with contribu;on from all the authors; Visualiza;on: PV, KS, FH, MB; Supervision: PV, CS, JR; Funding Acquisi;on: PV

## Resource availability

Further information and requests for resources and reagents should be directed to and will be fulfilled by the lead contact, Paola Vagnarelli

All unique/stable reagents generated in this study are available from the lead contact with a completed materials transfer agreement.

## Data and code availability

<u>The ATAC seq datasets</u> are deposited in arrayexpress and will be made publicly available upon publication. The reviewer can access the files using this link.

https://www.ebi.ac.uk/biostudies/arrayexpress/studies/E-MTAB-13826?key=4c721c35-81c8-42ba-a7fb-93aa2b030f46

E-MTAB-13826 Study typeATAC-seq EFO

<u>The RNA seq datasets</u> are deposited in arrayexpress and will be made publicly available upon publication. The reviewer can access the files using this link.

https://www.ebi.ac.uk/biostudies/arrayexpress/studies/E-MTAB-13827?key=8713847e-900b-4201-ab50-61efe98f75c0

AccessionE-MTAB-13827

Study typeRNA-seq of coding RNA EFO

<u>The Mass spectrometry dataset</u> are deposited in PRIDE and will be made publicly available upon publication. The reviewer can access the files using this link.

**Project Name:** Differential contribution of the chromatin-associated PP1 binding proteins PNUTS, Repo-Man and Ki-67 to mitotic exit and G1 re-establishment.

**Project accession:** PXD049319

**Password:** iVAd7u6R

## MATERIALS AND METHODS

### Experimental model and subject details

HCT116 cells were grown in McCoy’s 5A Medium GlutaMAX (Gibco) supplemented with 10% foetal bovine serum (Labtech), 100 U/mL penicillin and 100 µg/mL streptomycin (Gibco) in a humidified incubator at 37°C with 5% CO_2_.

HCT116-Ki-67-mAID-mClover-HygR/NeoR, HCT116-Ki-67-mAID-mClover-HygR/NeoR + DHB-mCherry and HCT116-Ki-67-mAID-mClover-HygR/NeoR + GEMININ-mCherry cell lines were generated in the Vagnarelli’s lab as previously described (DOI: https://doi.org/10.1101/2023.04.18.537310).

The HCT116 cell line stably expressing OsTIR1 under TET-ON promoter (HCT116-TET-OsTIR1) was kindly provided by Dr Masato T. Kanemaki (University of Tokyo, Japan).

## METHODS DETAILS

### Plasmids

pX330-U6-Chimeric_BB-CBh-hSpCas9 (no. 42230, Addgene) plasmids containing guide RNAs (gRNAs), designed with CRISPOR (http://crispor.tefor.net), were generated according to (Ran et al., 2013). The sequences of gRNAs are listed in Table S1.

To generate the plasmid donors containing Repo-Man or PNUTS homology arms, genomic DNA from HCT116-TET-OsTIR1 cells was purified by using EchoLUTION CellCulture DNA Kit (BioEcho), and the genomic DNA region around the stop codon of Repo-Man (1.2 kb) or PNUTS (1.3 kb) was amplified using Phusion High-Fidelity DNA Polymerase (Thermo Fisher Scientific; primers are listed at Table S1), cloned into pGEM-T Easy empty vector (Promega) and sequenced. Based on sequencing, plasmids containing Repo-Man or PNUTS homology arms were synthesized, with the following changes: the stop codon was mutated into a BamHI restriction site and silent point mutations were introduced within PAM sequences. The mAID-mClover-HygR (hygromycin resistance) cassette and mAID-mClover-NeoR (neomycin resistance) cassette (Natsume et al., 2016) were then inserted into the BamHI site, in frame with Repo-Man or PNUTS to generate homology arms donor plasmids.

### Cell line generation

For the generation of Repo-Man/PNUTS-mAID-mClover-HygR/NeoR cell lines, HCT116-TET-OsTIR1 cells were transfected with 2 µg of total DNA of pX330-U6-Chimeric_BB-CBh-hSpCas9 and homology arms plasmids (Repo-Man-mAID-mClover-HygR and Repo-Man-mAID-mClover-NeoR or PNUTS-mAID-mClover-HygR and PNUTS-mAID-mClover-NeoR for mAID cell lines; or -APEX2-mClover-HygR and -APEX2-mClover-NeoR for APEX cell lines) in 1:1:1 ratio, and selected for 14 days with 100 μg/mL hygromycin B (10687010, Invitrogen) and 700 μg/ml of geneticin (10131027, Gibco). Clones were first assessed by microscopy and then screened via PCR genotyping using 3-5 different combinations of primers, which targeted sequences outside of homology arms and within tagging inserts (Table S1).

For the generation of HCT116-Repo-Man/PNUTS-mAID-mClover-HygR/NeoR + DHB-mCherry stable cell lines, HCT116-Repo-Man/PNUTS-mAID-mClover-HygR/NeoR cells were co-transfected with 2 μg of total DNA of CSII-pEF-hDHB-mCherry and pMSCV-Blasticidin in 9:1 ratio and selected with 10 μg/ml blasticidin (15205, Merck). Clones were chosen by microscopy.

For the generation of HCT116-Ki-67/Repo-Man/PNUTS-mAID-mClover-HygR/NeoR + Geminin-mCherry stable cell lines, the same process was followed as the DHB-mCherry cell line generation.

### Mitotic synchronisation and protein degradation for G1 analyses

Unless stated otherwise, HCT116 cells (Ki-67 or Repo-Man or PNUTS-mAID-mClover-HygR/NeoR) were treated with 2 μg/ml doxycycline (D9891, Merck) and 0.06 μg/ml nocodazole (M1404, Merck), 24 and 18 h respectively, before the addition of 1 mM IAA (I2886, Merck) to induce degradation of mAID-tagged proteins. After 4 h treatment with IAA, mitotic cells were manually detached by shake-off, and cells were transferred to new poly-L-lysin coated dishes. Once attached, cells were washed 3x with PBS and released from mitosis into G1 for 18 h in a fresh medium containing 2 mM thymidine (T9250, Merck).

### Immunofluorescence microscopy

Cells were fixed with 4% PFA (10131580, Fisher Scientific) in PBS and processed as previously described (Vagnarelli et al., 2006). Primary and secondary antibodies used in the study are listed in Table S2. Three-dimensional data sets were acquired using a wide-field microscope (Delta Vision) Cascade II:512 camera system (Photometrics) and Olympus UPlanSApo 100x/1.40na Oil Objective (Olympus) and a wide-field microscope (NIKON Ti-E super research Live Cell imaging system) with a 100X Plan Apochromat lens, numerical aperture (NA) 1.45. The data sets were deconvolved using Delta Vision software. Three-dimensional data sets were converted to a maximum projection, exported as PNG files and imported into Inkscape software (version 0.91) for final presentation.

For deepSIM imaging a NIKON Ti2-E inverted microscope with 100X Plan Apochromat lens, numerical aperture (NA) 1.45 with a sCMOS camera.3D stacks were acquired and reconstructed with the NIS software.

### Immunoblotting

Whole-cell extracts were prepared by direct lysis in 1x Laemmli sample buffer (Laemmli, 1970).

For histone isolation, HCT116 cells were synchronised, proteins of interest were degraded as described above and collected at the G1 stage of the cell cycle. 5×10^6^ cells were collected from each condition and incubated for 10 min on ice with 500 µl lysis buffer (1 mM Tris-HCl, pH 7.5, 10 mM NaCl, 0.5% NP-40) supplemented with protease and phosphatase inhibitors. After 3 min centrifugation at 1,000 *g* at 4°C, the supernatant was discarded, the pellet was resuspended in 500 µl lysis buffer and samples were pelleted again as above. The pellet was flash-frozen in liquid nitrogen and stored at -20°C for 1 h. Next, the pellet was thawed on ice for 15-30 min and washed with 400 µl of digestion buffer (50 mM Tris-HCl, pH 7.5, 1 mM CaCl_2_, 4 mM MgCl_2_, 300 mM sucrose). After 10 min centrifugation at 100 *g* at 4°C, the supernatant was discarded, and the pellet was resuspended in 300 µl of digestion buffer with 600 U of micrococcal nuclease (MNase) (M0247, New England Biolabs) and incubated for 30 min at 37°C. The MNase reaction was terminated by adding 1mM EDTA. The digested sample was then centrifugation for 20 min at 8,000 *g* at room temperature (RT) and the pellet was resuspended in 1x Laemmli sample buffer.

The proteins were separated by SDS–PAGE and transferred onto nitrocellulose membranes (10600003, Amersham). Membranes were blotted with primary and secondary antibodies, which are listed in Table S2, and visualised using the LiCor Odyssey system or g-Box Chemi Xrq Gel Doc System.

### CDK2 activity analyses

HCT116 cell lines containing mAID-tagged proteins and stably-expressed DHB-mCherry were synchronised, and proteins were degraded as described above. Upon reaching the G1 stage of the cell cycle, cells were fixed with 4% PFA in PBS. Coverslips were mounted with VECTASHIELD® Antifade Mounting Medium (H-1000-10, Vector Laboratories) containing DAPI, and observed on a wide-field microscope (NIKON Ti-E super research Live Cell imaging System). The analysis of CDK2 activity was conducted in ImageJ according to (Cappell et al., 2016). Violin plots were generated with R Studio. The Wilcoxon statistical analysis was conducted with R Studio.

### APC/C activity analyses

HCT116 cell lines containing mAID-tagged proteins and stably-expressed Geminin-mCherry were treated as DHB-mCherry-expressing cells described above. The analysis of APC/C activity was conducted in Fiji according to (Cappell *et al*., 2016). Violin plots were generated with R Studio. The Wilcoxon statistical analysis was conducted with R Studio.

### Comet assay

The alkaline comet assay was adapted from (Tice et al., 2000). Poly-L-lysine coated slides were first covered with 1% high-melting-point agarose and dried overnight at room temperature. After cell treatment, a drop of normal-melting-point agarose was first loaded on a slide, and then a drop of low-melting-point agarose was put on the precoated slide. Then, 10^4^ cells were placed in the agarose droplet. Slides were lysed in a detergent solution (containing 2% N-Lauroylsarcosine sodium salt, 0.5 M Na_2_EDTA and 0.1 mg/ml proteinase K), for 1 hour at 4°C. DNA unwinding was carried out with a neutral solution at pH 8.5 (90 mM Tris-HCl, 90 mM boric acid and 2 mM Na_2_EDTA) for 90 minutes at RT in a 1-L electrophoresis unit. Electrophoresis was conducted for 40 minutes (20 V) in the same buffer solution at 4°C. After the electrophoretic run, the slides were neutralized with distilled water, stained with SYBR Safe in TE buffer, dehydrated by dipping into 70%, 90% and 100% ethanol, sequentially, and then dried overnight at RT. Images were captured using Leica DM4000 fluorescence microscope (Leica Microsystems). Comet length was measured in ImageJ.

### Fluorescence in situ hybridisation (FISH)

HCT116 cells with mAID-tagged proteins were synchronised and protein degraded as described above. Cells were fixed twice as follows: 15 min with 75 mM KCl, then 30 min with 3:1 ice-cold methanol:acetic acid. Finally, cells were stored in methanol/acetic acid at -20°C. P1-derived artificial chromosome (PAC)-derived probes were extracted from bacterial cultures and fluorescently labelled by nick translation using Nick Translation Kit (Abbott), following the manufacturer’s instructions. The probe was separated using G-50 column, precipitated at - 20°C in the presence of 10x salmon sperm DNA for the centromeric probes, according (Garimberti and Tosi, 2010).

### Pericentromeric/ α-satellite FISH

The probe (3 µl of probe and 7 µl of hybridisation buffer: 50% (v/v) formamide, 10% (w/v) dextran sulphate, 1x Denhart’s, 2x SSC, pH 7.0) was denatured at 73°C for 5 min. Probe and nuclei were denatured at 85°C for 5 min, and then hybridised overnight at 39°C. Slides were washed with 2x SSC at RT for 10 min, 0.4x SSC at 60°C for 10 min, 0.1x SSC at RT for 10 min and counterstained with DAPI.

For all FISH analyses, three-dimensional data sets were acquired using a wide-field microscope (NIKON Ti-E super research Live Cell imaging system) with 100x, 1.45 (NA) Plan Apochromat lens. The data sets were deconvolved with NIS Elements AR analysis software (NIKON). Three-dimensional data sets were converted to maximum projection in the NIS software, exported as TIFF files, and imported into Inkscape software (version 0.91) for final presentation.

### Proximity ligation assay (PLA)

Proximity ligation assay was performed according to the manufacturer’s protocol (Sigma). HCT116:Ki-67-AID cells were treated as indicated in Figure 1A or with Doxycycline (2μg/ml) and 0.06 μg/ml Nocodazole for 24 and 18 h before the addition of Auxin 1000μM f, respectively. After the completion of the experiment indicated in Figure 1 A or 4 h treatment with Auxin the cells were fixed, permeabilized and blocked with BSA as previously described [65]. The antibodies were used at a concentration as follows, 1:1000 anti-CENP-B (gift from W.C. Earnshaw, Edinburgh) and 1:10,000 anti-GFP [PABG1] (Cat# PABG1-20, RRID:AB_2749857). PLA probes were added, and ligation was performed following the manufacturer instructions (Sigma). Coverslips were mounted with vectashield containing DAPI and observed on the previously mentioned wide-field NIKON microscope.

### RNA sequencing

Total RNA was extracted from HCT116 cells, with or without the protein of interest following the experiment described in Figure1A, using Monarch Total RNA Miniprep Kit (New England Biolabs), according to manufacturer’s instructions. RNA samples were sent to Macrogen Europe B.V (The Netherlands). Macrogen Europe BV constructed libraries using Illumina TruSeq stranded mRNA library preparation with Ribozero rRNA depletion. Sequencing was performed with a Novaseq 6000 platform, at 100 M paired-end reads per sample.

The trimmed reads were aligned to the human reference genome GRCh38, using HISAT2 (v2.2.1) under standard conditions.

The resulting alignments were filtered for high quality hits using SAMtools v0.1.12 (Li et al., 2009) with a minimum selection threshold score of 30. FeatureCounts function (from Subread package v2.0.2) was used to count gene level reads, and Deseq2 was used to identify differentially expressed genes between samples. The differential expression was expressed in the form of log2 fold change between sample and control and deemed statistically significant by a lower p-value of 0.05.

Functional enrichment was analysed using STRING (string-db.org), while Venn diagrams were performed in the free software FunRich. Volcano plots were performed using the ggplot package (v3.4.2) in R Studio.

### ATAC sequencing

Synchronised HCT116:RIPPOs-AID cells, with or without IAA were used following the experiment described in Figure1 A. For each cell line, two replicates containing 100,000 cells per condition were submitted to Active Motif for ATAC-sequencing.

Alignment of the ATAC-seq data was performed against the human reference genome GRCh38. Peaks calling was made using MACS 2.1.0 with 1e-7 as cut-off of p-value, without control file and with the -nomodel parameter. Peaks were filtered using the ENCODE blacklisted regions. FRIP (fraction of reads in peaks) value for each sample is higher than 25%.

Bioconductor (v3.16) packages DiffBind and DESeq2 were used to identified differentially accessible regions between samples.

ChIP-seq datasets for H3K9ac and H3K27ac were downloaded from the public database ENCODE (ENCFF105NDA and ENCFF853VVI, respectively). Overlaps between the differentially accessible region identified in our ATAC-seq dataset and H3K9ac or H3K27ac ChIP-seq datasets were identified with bedtools 2.31.0. Statistical significance of these overlaps was given by Fisher’s test (fisher function with bedtools).

Hg38 Repeat file was downloaded from UCSC Genome browser (https://genome.ucsc.edu) and overlaps between the differentially accessible region identified in our ATAC-seq dataset and the repeat dataset was made using bedtools 2.31.0. Statistical significance of these overlaps was given by Fisher’s test (fisher function with bedtools).

### Mass spectrometry

Protein samples were processed at the same time and using the same digestion protocol without any deviations. They were subjected for MS analysis under the same conditions. Protein and peptide lists generated using the same software and the same parameters. Specifically, 30 µg of total protein from each sample were digested using the Filter Aided Sample Preparation (FASP) protocol as described by (Wisniewski et al., 2009) with minor modifications. In brief, each protein sample was added on the top of a 30 kDa MWCO filter units (Vivacon, UK) along with 150 µl of denaturation buffer (8M Urea in 50mM ammonium bicarbonate (ABC) (Sigma Aldrich) and spun at 14,000 x g for 20 min, while another wash with 200 µl of denaturation buffer was performed under the same conditions. The protein samples were then reduced by the addition of 100 µl of 10 mM dithiothreitol (Sigma Aldrich, UK) in denaturation buffer for 30 min at ambient temperature, and alkylated by adding 100 µl of 55 mM iodoacetamide (Sigma Aldrich, UK) in denaturation buffer for 20 min at ambient temperature in the dark. Two washes with 100 µl of denaturation buffer and two with digestion buffer (50mM ABC) were performed under the same conditions described above before the addition of trypsin (Pierce, UK). The protease:protein ratio was 1:50 and proteins were digested overnight at 37°C. Following digestion, samples were spun at 14,000 x g for 20 min and the flow-through containing digested peptides was collected. Filters were then washed one more time with 100 µl of ABC and the flow-through was collected again. The eluates from the filter units were acidified using 20 µl of 10% Trifluoroacetic Acid (TFA) (Sigma Aldrich, UK), and 2% of the eluate was spun onto StageTips as described by (Rappsilber et al., 2007). The rest 98% of the eluate was subjected to phospho-enrichment using Ti-IMAC MagResyn® beads (ReSyn Biosciences), following the beads’ protocol without deviations. Following enrichment, peptides were concentrated and cleaned using the same StageTip protocol as above.

In both cases, peptides were eluted in 40 μL of 80% acetonitrile in 0.1% TFA and concentrated down to 1 μL by vacuum centrifugation (Concentrator 5301, Eppendorf, UK). The peptide sample was then prepared for LC-MS/MS analysis by diluting it to 5 μL by 0.1% TFA.

LC-MS analyses were performed on an Orbitrap Exploris™ 480 Mass Spectrometer (Thermo Fisher Scientific, UK) coupled on-line, to an Ultimate 3000 HPLC (Dionex, Thermo Fisher Scientific, UK). Peptides were separated on a 50 cm (2 µm particle size) EASY-Spray column (Thermo Scientific, UK), which was assembled on an EASY-Spray source (Thermo Scientific, UK) and operated constantly at 50oC. Mobile phase A consisted of 0.1% formic acid in LC-MS grade water and mobile phase B consisted of 80% acetonitrile and 0.1% formic acid. Peptides were loaded onto the column at a flow rate of 0.3 μL min-1 and eluted at a flow rate of 0.25 μL min-1 according to the following gradient: 2 to 40% mobile phase B in 120 min and then to 95% in 11 min. Mobile phase B was retained at 95% for 5 min and returned back to 2% a minute after until the end of the run (160 min).

Survey scans were recorded at 120,000 resolution (scan range 350-1500 m/z) with an ion target of 3.0e6, and injection time of 50ms and RF lens of 40%. MS2 was performed in the orbitrap in Data Dependent Acquisition (DDA) mode at 15,000 resolution with isolation window of 1.4, maximum injection time of 50ms and AGC target of 8.0E5 ions. We used HCD fragmentation (Olsen et al., 2007) with stepped collision energy of 30. Data for both survey and MS/MS scans were acquired in profile mode.

The MaxQuant software platform (Cox and Mann, 2008) version 1.6.1.0 was used to process the raw files from the SILAC labelled samples and the search was conducted against the complete/reference proteome set of Homo sapiens (Uniprot database - released in 2019), using the Andromeda search engine (Cox et al., 2011). For the first search, peptide tolerance was set to 20 ppm while for the main search peptide tolerance was set to 4.5 pm. Isotope mass tolerance was 2 ppm and maximum charge to 7. Digestion mode was set to specific with trypsin allowing maximum of two missed cleavages. Carbamidomethylation of cysteine was set as fixed modification. Oxidation of methionine and phosphorylation of serine, threonine and tyrosine were set as variable modifications. To account for the SILAC labels, multiplicity was set to 2, with lysine +8 and arginine +10 selected. FDR was set to 1%.

## QUANTIFICATION AND STATISTICAL ANALYSES

### FISH analyses

For the signal area analyses, 3D stack images were exported and analysed with ImageJ. Scale was set to 1 μm, ROIs of the signals were created using threshold function and wand tracing tool, the area in μm was used to generate the violin plots.

### CENP-B

Quantification of the kinetochore/centromere staining. CENP-B, 3D stack images of control and Ki-67-degraded cells; were exported and analysed with the Foci quantification plug-in using Fiji (1_color_auto.ijm) (Ledesma-Fernandez and Thorpe, 2015). Background was subtracted, and the mean intensity was used to generate the violin plots.

### PLA

Spots laying within nuclear masks were counted in control and Ki-67-degraded cells; the numbers of foci were used to generate the violin plots.

### Nuclear area

For the signal area analyses, 3D stack images were exported and analysed with ImageJ. Scale was set to 1 μm, ROIs of the nuclear area were created, the area in μm was used to generate the violin plots.

### RNA Polymerase II (POL II)

For each HCT116 cell line (Ki-67 or Repo-Man or PNUTS), images were acquired for three biological replicates using a wide-field microscope (NIKON Ti-E super research Live Cell imaging system) with 40x, 1.45 (NA) Plan Apochromat lens. Analysis was then conducted in NIS Elements AR analysis software (NIKON). Using nuclei as a mask, fluorescent intensities of RNA POL II and phospho-S5-RNA POL II were measured. Background of each channel from each image w subtracted and the ratio of fluorescence intensity of phospho-S5-RNA POL II/total RNA POL II was used to generate the violin plots

### Statistical analyses

Statistical analyses were performed either in Excel (chi-squared test) or in R (using the Wilcoxon rank test function, differential expression, lowest smoothing).

**Table S1:**
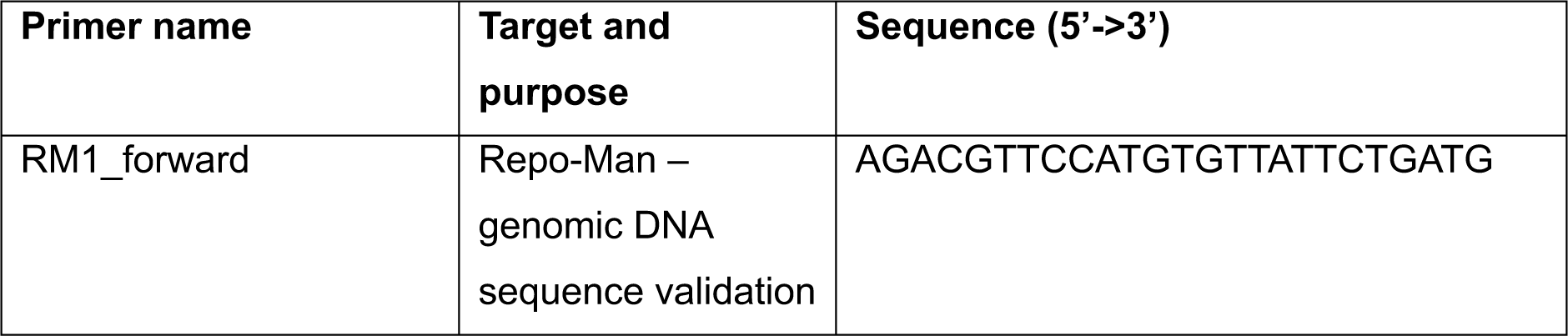
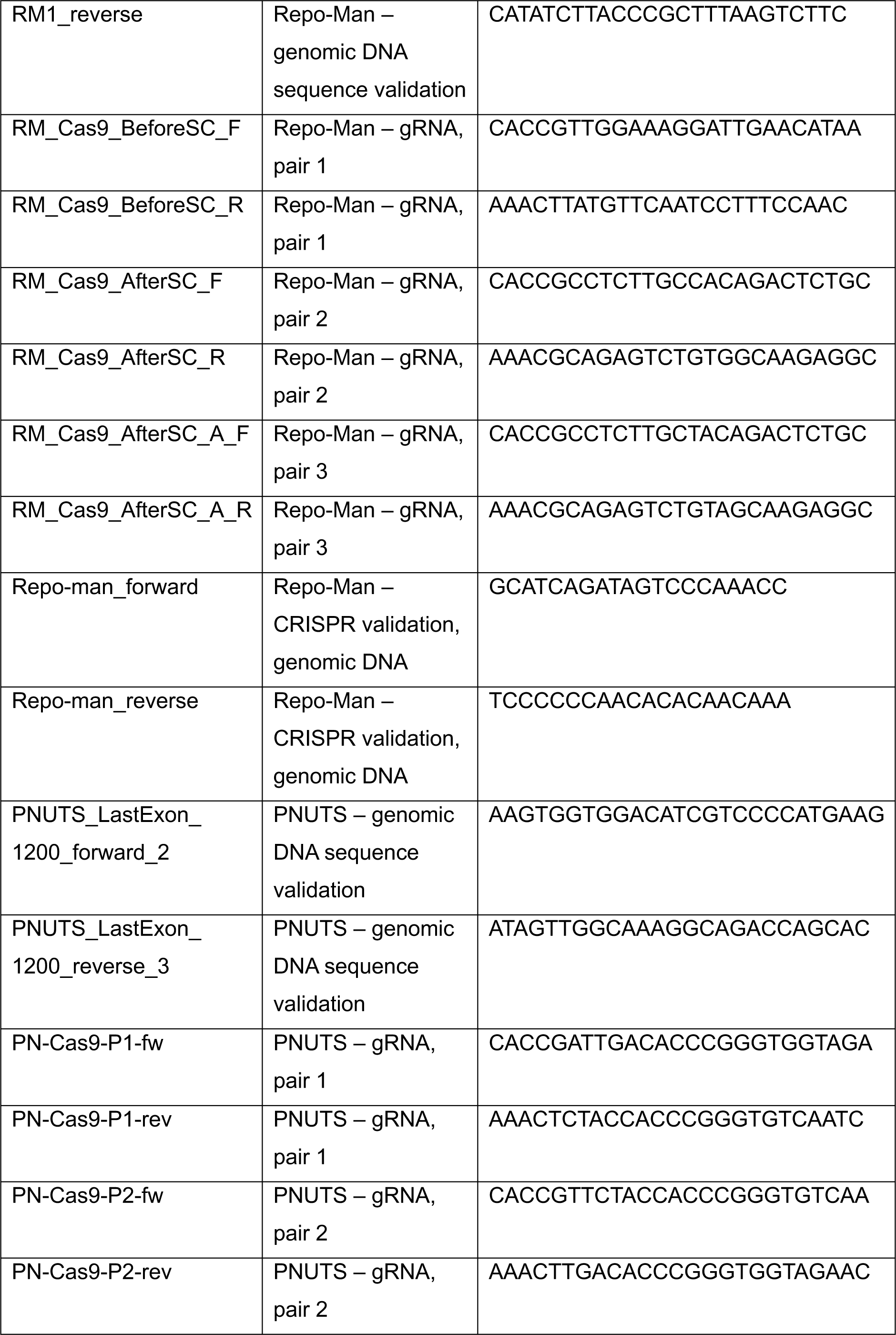
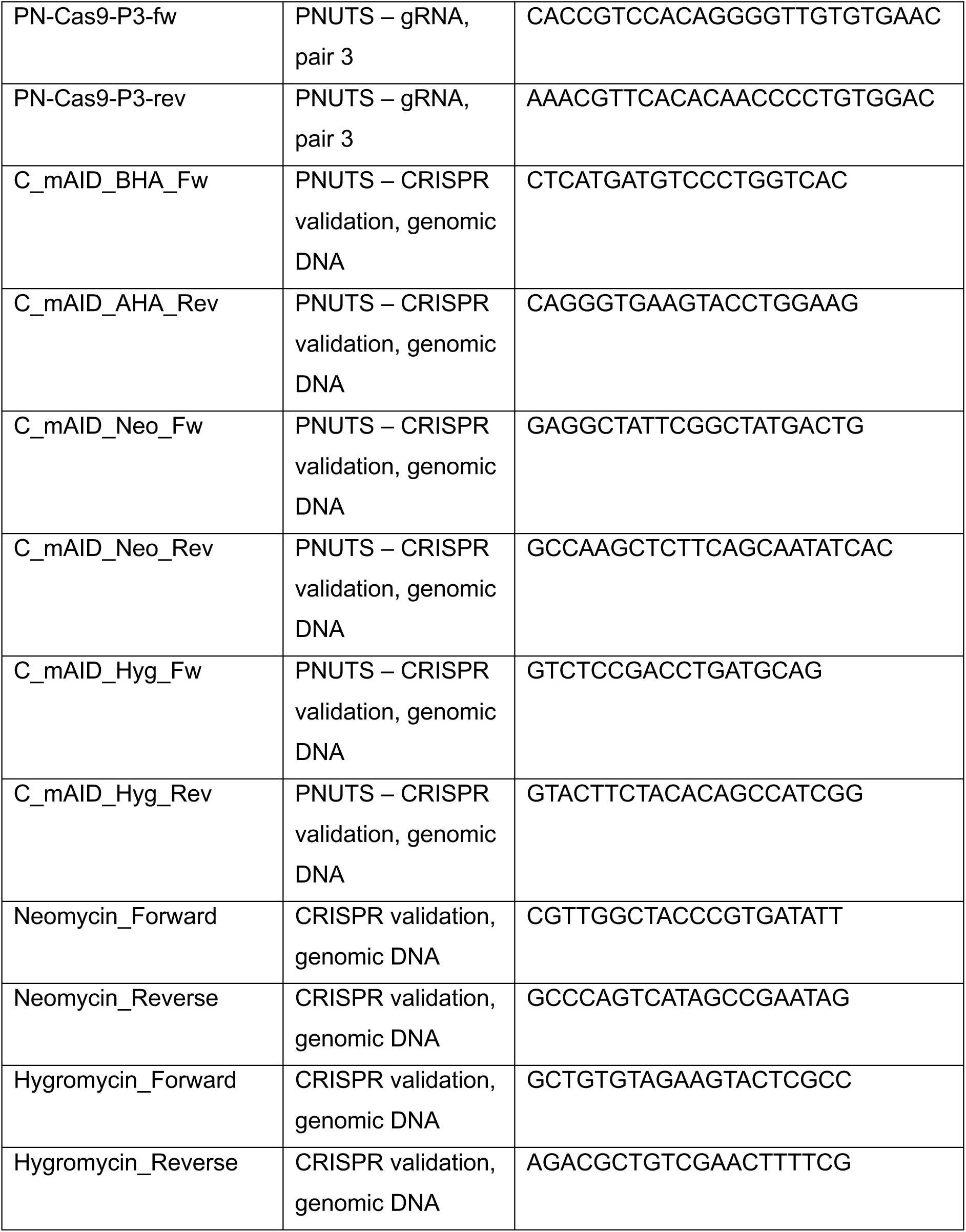
List of primers used in the study.

**Table S2:** List of antibodies used in the study.

| Target | Dilution<br>(ICC/WB)* | Species | Catalogue<br>number | Supplier |
| --- | --- | --- | --- | --- |
| α-tubulin (B-5-1-2) | 1:1,000 (ICC) | mouse | T5168 | Sigma Aldrich |
| dsDNA | 1:1,000 (WB) | mouse | MAB030 | Millipore |
| GAPDH | 1:5,000 (WB) | mouse |  | Proteintech |
| H3K9ac | 1:1000 (WB) | rabbit | ab10812 | Abcam |
| H3K9me | 1:1000 (WB) | mouse |  | Active Motif |
| H3K27ac | 1:1000 (WB) | rabbit | 39133 | Active Motif |
| H3K27me2/3 | 1:1000 (WB) | mouse | 39535 | Active Motif |
| H4K20me1 | 1:1000 (WB) | mouse | 39027 | Active Motif |
| Lamin A/C | 1:1,000 (ICC) | rabbit | ab108595 | Abcam |
| Retinoblastoma (Rb) | 1:250 (ICC) | mouse | 554136 | BD Biosciences |
| Retinoblastoma (phospho-Rb) | 1:250 (ICC) | rabbit | 8516 | Cell Signaling |
| Repo-Man | 1:500 (WB) | Rabbit | ab45129 | Abcam |
| RNA Polymerase II | 1:1,000 (ICC) | mouse | sc-55492 | Santa Cruz |
| RNA Polymerase II (phospho-S5) | 1:1,000 (ICC) | rabbit | ab5131 | Abcam |
| R-loops (S9.6) | 1:1,000 (WB) | mouse | MABE1095 | Millipore |
| CENP-B | 1:1,000 (IF) | mouse |  | Earnshaw Lab |

\*Abbreviations: *ICC*, immunocytochemistry; *WB*, western blotting.

## FIGURE LEGENDS

**Supplementary Figure 1.**
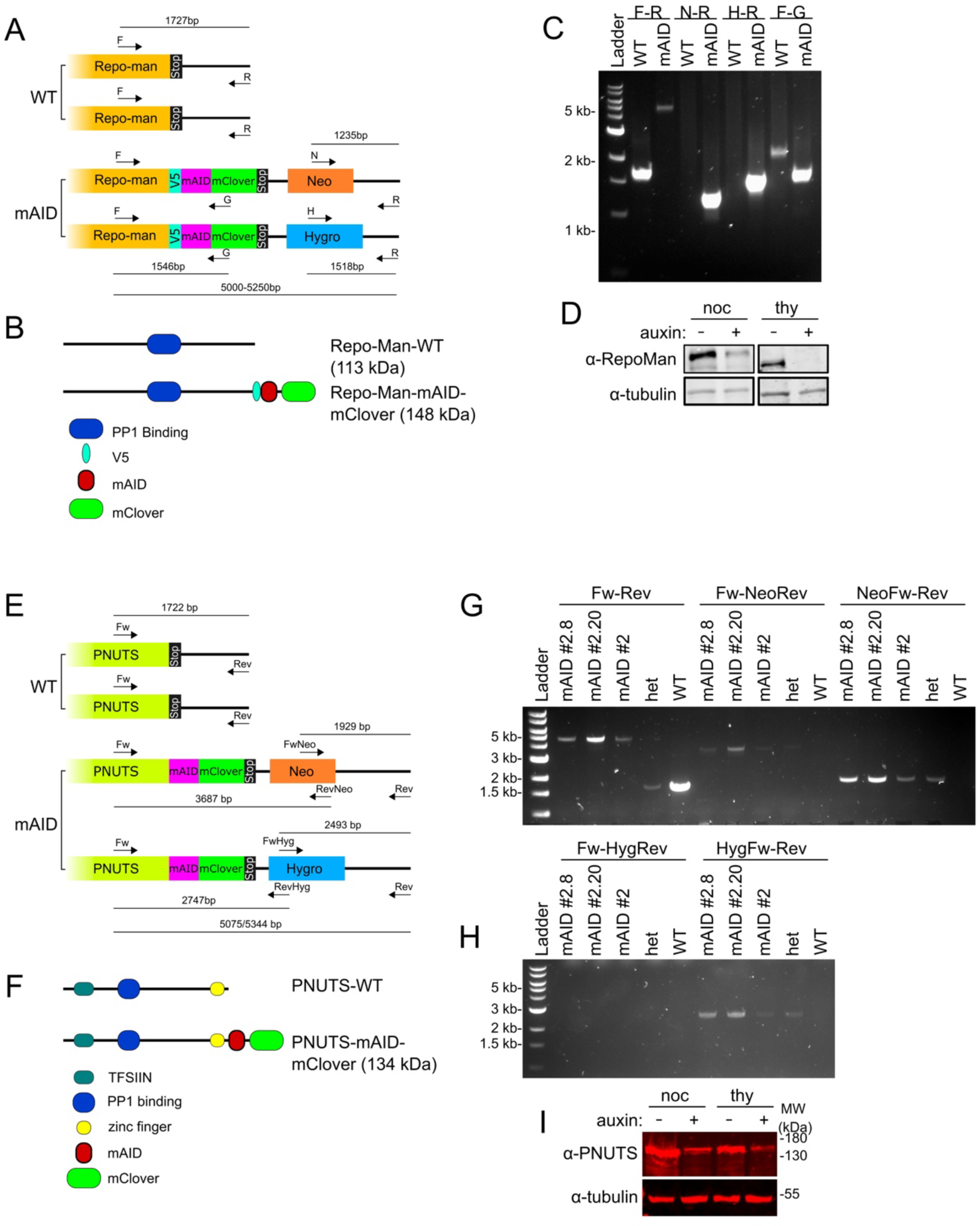
A, B, E, F) Schematic representation of CRISPR constructs designed to target both alleles of Repo-Man (A) and PNUTS (E), and PCR genotyping of the endogenously tagged AID cell lines, HCT116:RepoMan-mAID (B) and HCT116:PNUTS-mAID (F). Several primer pairs (as denoted in A and E), targeting regions inside and outside of homology arms, were used for validation. M=1kb ladder C, D, G, H) Schematic representation of Repo-Man (C) and PNUTS (G) wild-type and predicted new proteins with domains of interest. Western blots of whole cell lysate of HCT116:RepoMan-mAID (D) and HCT116:PNUTS-mAID (H) cell lines treated with IAA during nocodazole-mediated mitotic synchronisation and release into G1 in the presence of thymidine. The blots were probed with anti Repo-Man (D, top panels), anti PNUTS (H, top panels) and anti alpha tubulin (D and H, bottom panels) antibodies.

**Supplementary Figure 2.**
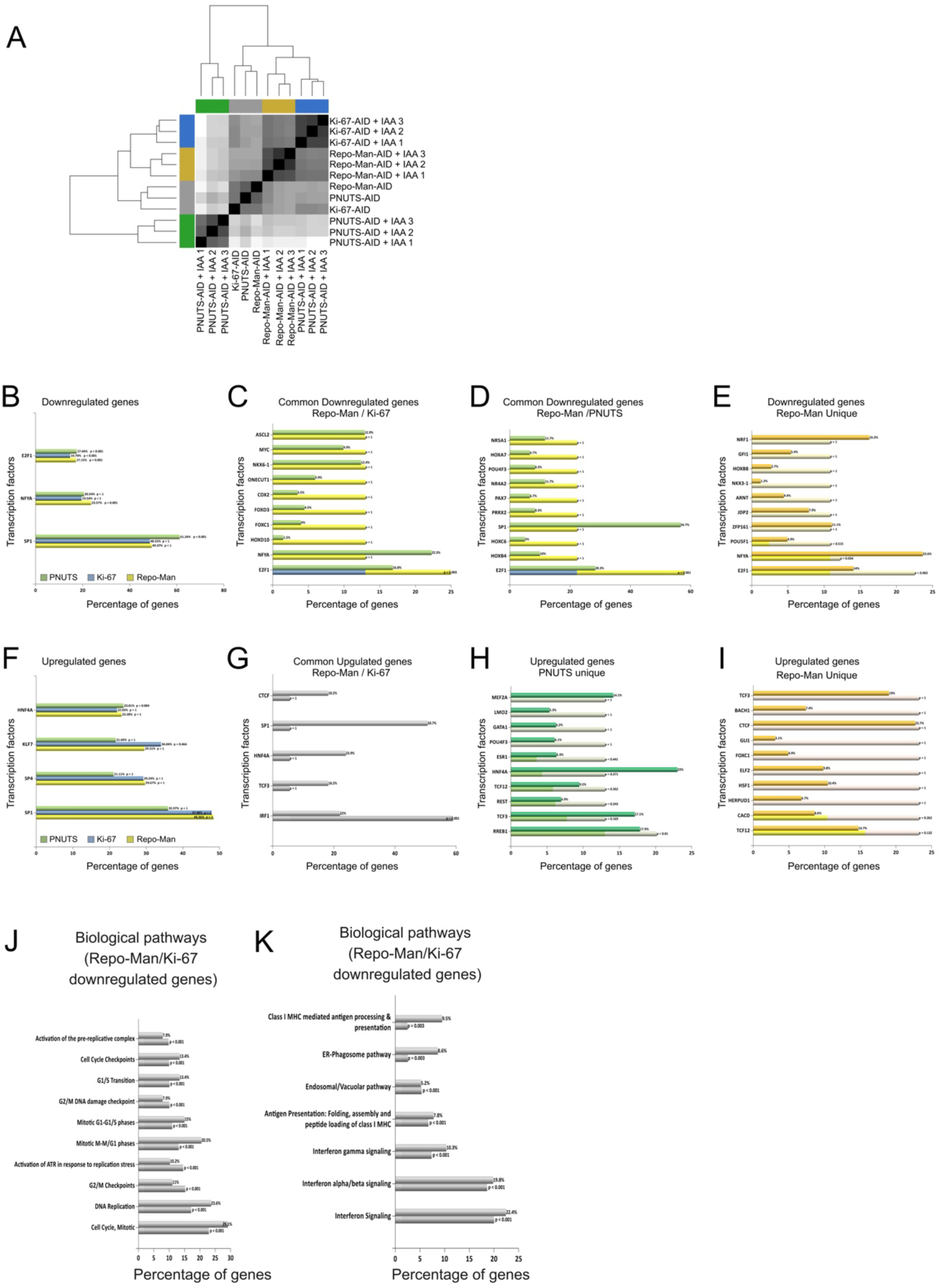
A) Heatmap of the distance matrix from the 3 control cell lines used in the RNA-seq analysis. B, C, D, E) FunRich transcription factors enrichment analysis of the downregulated genes common between Ki-67, Repo-Man and PNUTS (B), Repo-Man and Ki-67 (C), Repo-Man and PNUTS (D) or unique to Repo-Man (E). Downregulated genes from RNA-seq experiments of the HCT116:Ki-67-AID, HCT116:RepoMan-mAID and HCT116:PNUTS-mAID cell lines in Figure 1 E,F and G with and without IAA. F, G, H, I) FunRich transcription factors enrichment analysis of the upregulated genes common between Ki-67, Repo-Man and PNUTS (B), Repo-Man and Ki-67 (C), unique to PNUTS (D) or unique to Repo-Man (E). Upregulated genes were obtained by RNA-seq of the HCT116:Ki-67-AID, HCT116:RepoMan-mAID and HCT116:PNUTS-mAID cell lines in Figure 1 E,F and G. J, K) FunRich biological pathways enrichment analysis of the downregulated common (I) and upregulated common (L) genes between HCT116:Ki-67-AID and HCT116:RepoMan-mAID cell lines, obtained by RNA-seq in Figure 1 E and F.

**Supplementary figure 3.**
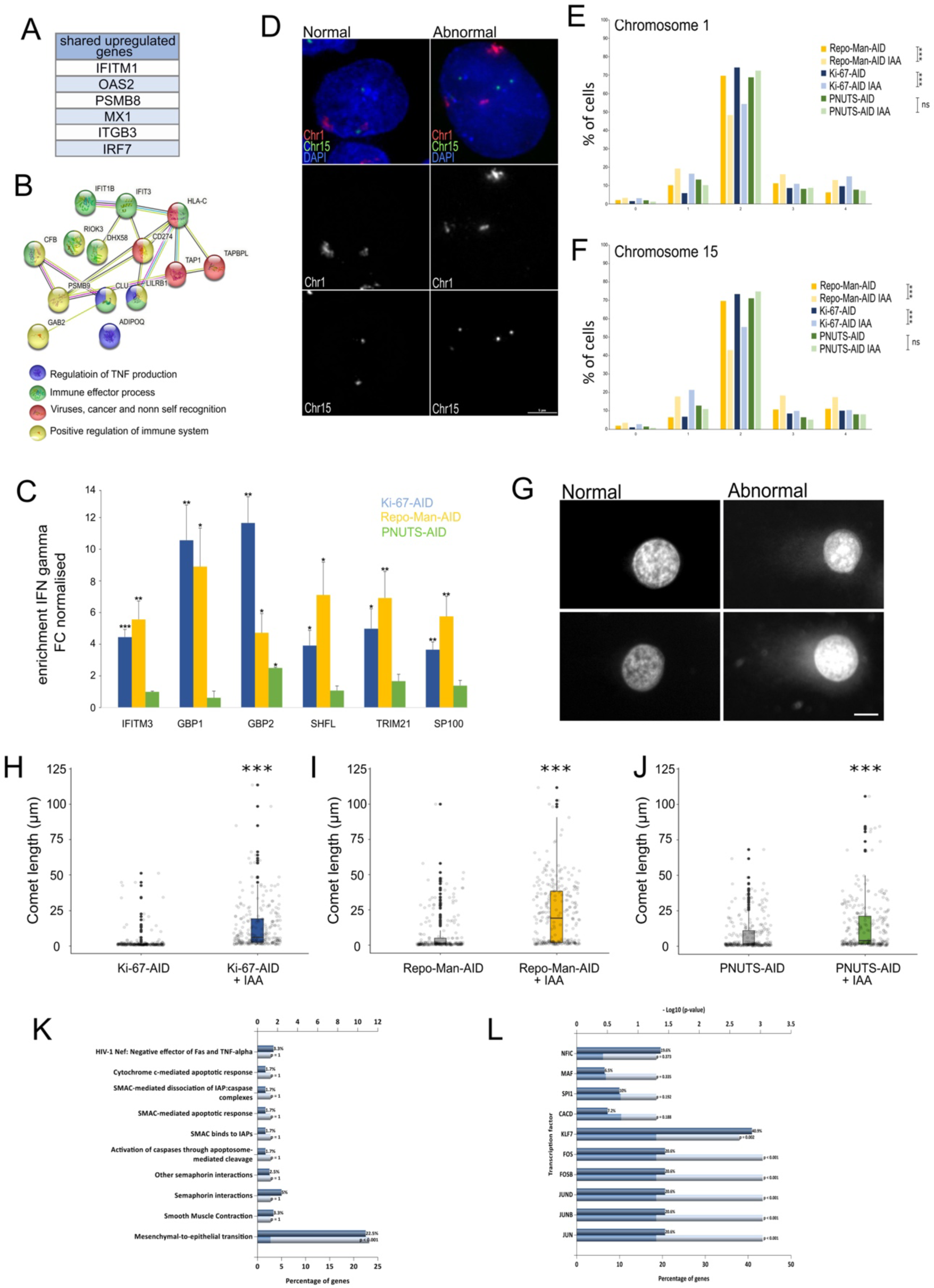
A) Common interferon genes between HCT116:Ki-67-AID, HCT116:RepoMan-AID and HCT116:PNUTS-AID cell lines obtained by RNA-seq in Figure 1 E, F and G. B) String pathway enrichment analyses of the common interferon genes between HCT116:Ki-67-AID and HCT116:RepoMan-AID cell lines obtained by RNA-seq in Figure 1 E and F. C) Enrichment of IFN gamma genes obtained from the RNA seq experiments described in Figure 1 E,F and G. The values represent the average of the 3 independent replicas and the error bars are the standard deviations. The experiments were analysed by a Student’s t-test. *= p<0.05 D) HCT116:Ki-67-AID, HCT116:RepoMan-AID and HCT116:PNUTS-AID cell lines were treated as indicated in Figure 1A, fixed and subjected to FISH. Representative images of the FISH normal (left) and abnormal (right) signals obtained with the pUC 177 (red signals, Chr 1) and pTRA-20 (green signals, Chr 15) probes. Scale bar 5 μm. E and F) Analysis of the foci patterns (0, 1, 2, 3 and 4 signals) of the experiment in (D) for Chromosome 1 (E) and Chromosome 15 (F). Sample size: CHR1: HCT116:Ki-67-AID: Control=528, IAA=535, HCT116:RepoMan-AID: Control=545, IAA=515 and HCT116:PNUTS-AID: Control=513, IAA=527, and CHR15: HCT116:Ki-67-AID: Control=554, IAA=582, HCT116:RepoMan-AID: Control=563, IAA=525 and HCT116:PNUTS-AID: Control=555, IAA=538. The data were statistically analysed with a Chi-squared test. ***= p<0.001, ns= not significant. G) Representative images of the comet assay of the HCT116:Ki-67-AID, HCT116:RepoMan-AID and HCT116:PNUTS-AID cell lines treated as indicated in Figure 1A. H, I and J) Quantification of the comet length of HCT116:Ki-67-AID (H), HCT116:RepoMan-AID (I) and HCT116:PNUTS-AID (J) cells lines. The box plots with jittered data represent the distribution of the comet tail length in μm. The box inside represents the 75th and 25th percentile, whiskers are the upper and lower adjacent values and the line is the median. A Wilcoxon test was conducted to compare the experiments. Sample size: HCT116:Ki-67-AID: Control=251, IAA=251, HCT116:RepoMan-AID: Control=225, IAA=273 and HCT116:PNUTS-AID: Control=263, IAA=275. *** = p<0.001 K) FunRich biological pathways enrichment analysis of the upregulated Ki-67 unique genes obtained by RNA-seq of the HCT116:Ki-67-AID cell line in the Figure 1 E. L) FunRich transcription factors enrichment analysis of the upregulated Ki-67 unique genes obtained by RNA-seq of the HCT116:Ki-67-AID cell line in the Figure 1 E.

**Supplementary figure 4.**
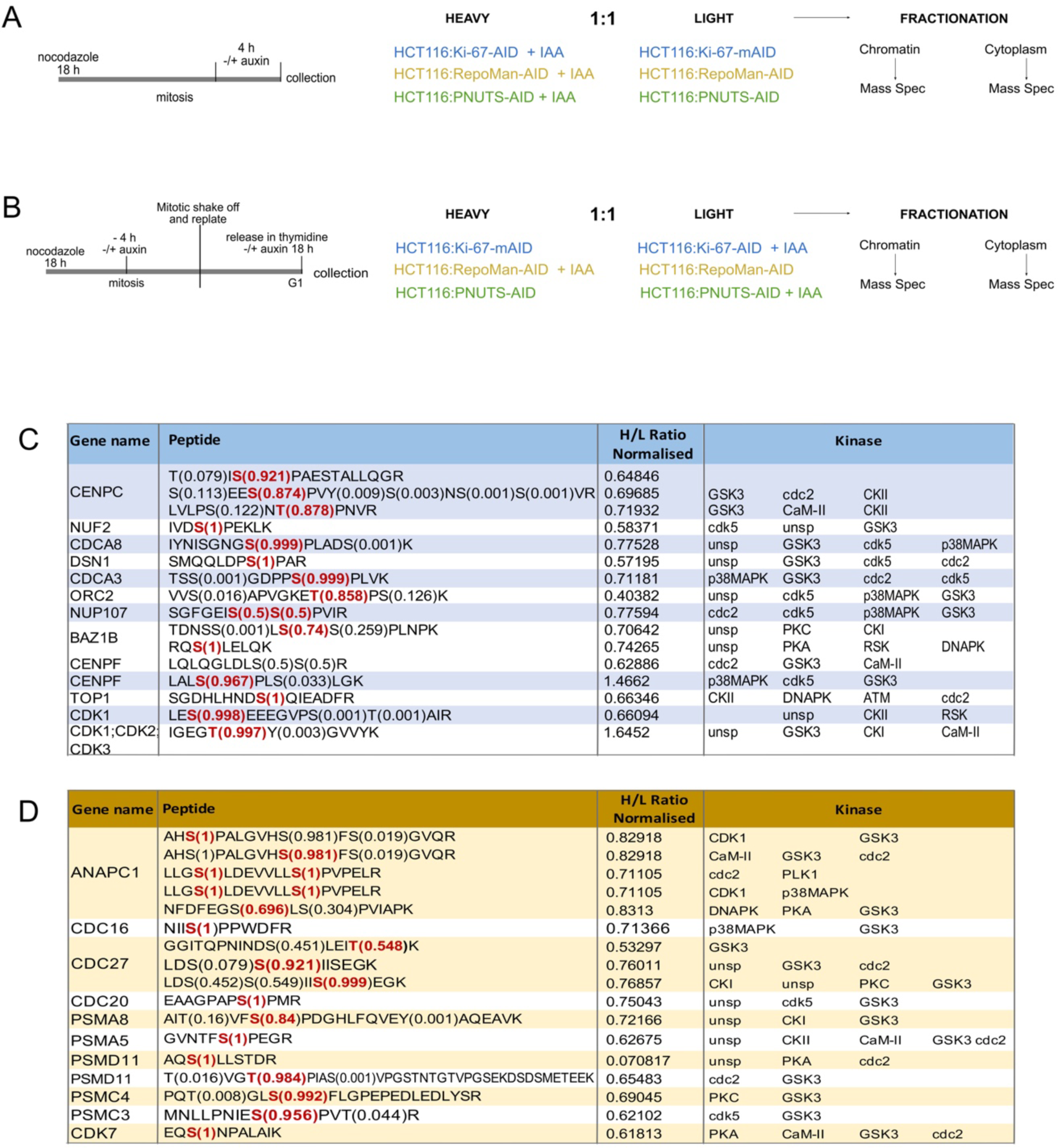
A) Scheme of the SILAC-based phospho-proteomic experiment in Mitosis. Samples collected for phosphoproteomic analyses from HCT116:Ki-67-AID, HCT116:RepoMan-AID and HCT116:PNUTS-AID cell lines treated with or without IAA for 4 h after 18 h incubation with nocodazole. Control samples were collected from cell lines cultured with light-labelled amino acids, whilst IAA samples were collected from cell lines cultured with heavy-labelled amino acids. B) Scheme of the SILAC-based phospho-proteomic experiment in G1. Samples collected for phosphoproteomic analyses from HCT116:Ki-67-AID, HCT116:RepoMan-AID and HCT116:PNUTS-AID following the experiment indicated in Figure 1A. Control samples were collected from HCT116:Ki-67-AID and HCT116:PNUTS-AID cell lines cultured with heavy-labelled amino acids, whilst IAA samples were collected from HCT116:Ki-67-AID and HCT116:PNUTS-AID cell lines cultured with light-labelled amino acids. Control samples were collected from HCT116:RepoMan-AID cell line cultured with light-labelled amino acids, whilst IAA samples were collected from HCT116:RepoMan-AID cell line cultured with heavy-labelled amino acids. C) Table of proteins and the differentially phosphorylated residues presented in Figure 6 A. In red highlighted the residues differentially phosphorylated and the phosphorylation probability. NetPhos - 3.1 was used for the prediction of the kinases regulating these phosphorylations. D) Table of proteins and the differentially phosphorylated residues presented in Figure 7 A. In red highlighted the residues differentially phosphorylated and the phosphorylation probability. NetPhos - 3.1 was used for the prediction of the kinases regulating these phosphorylations.

**Supplementary figure 5.**
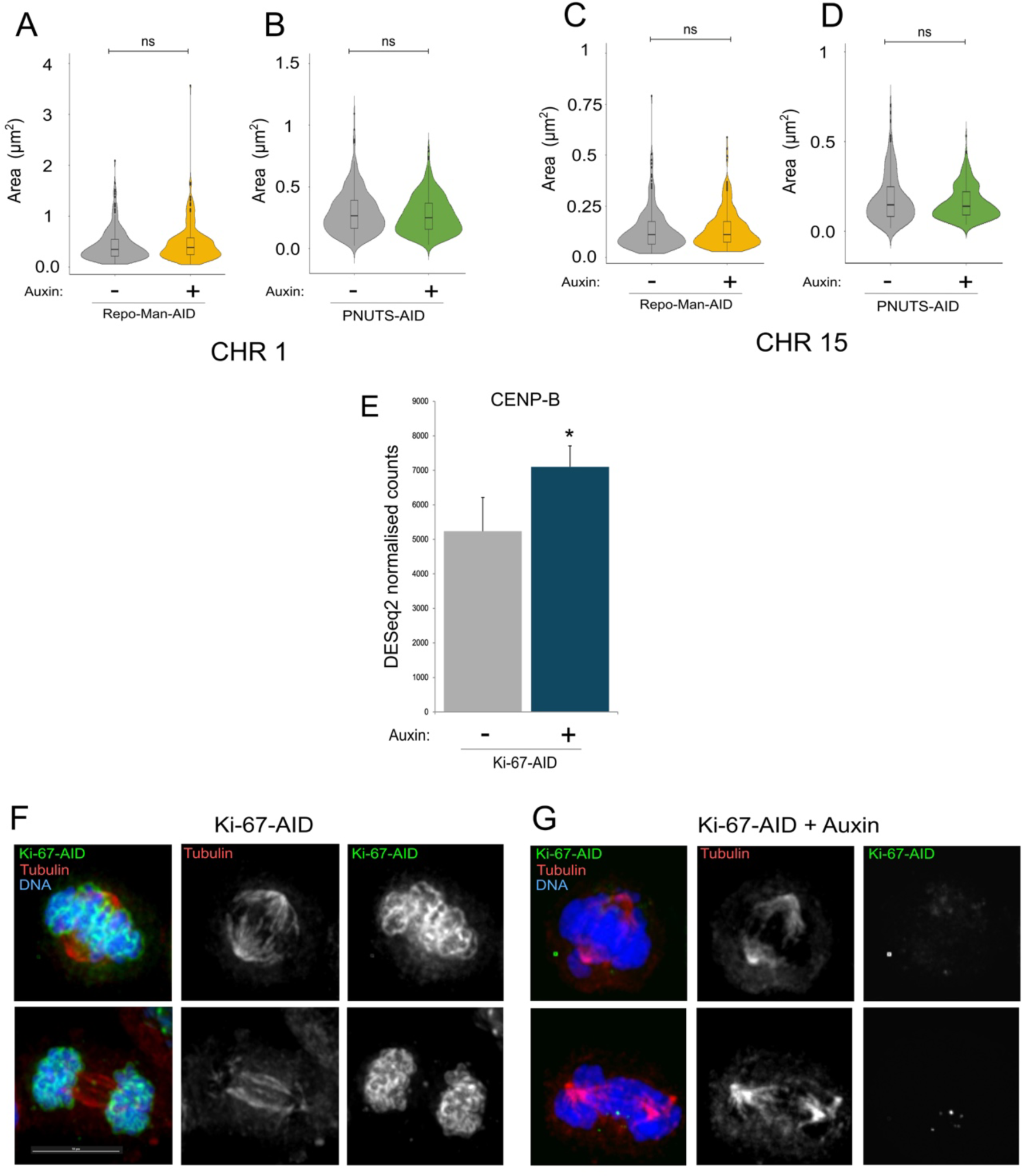
A, and B) HCT116:RepoMan-AID (A) and HCT116:PNUTS-AID (B) cell lines treated as in Figure 1 A fixed and subjected to FISH using pUC 177 probe. Quantification of the area occupied by the FISH signals. Sample size: HCT116:RepoMan-AID: Control=267, IAA=276, HCT116:PNUTS-AID: Control=346, IAA=405.The box inside the violin represents the 75th and 25th percentile, whiskers indicate the upper and lower adjacent values and the line is the median. The data presented in the violin plots were statistically analysed with a Wilcoxon test. ns= not significant. C and D) HCT116:RepoMan-AID (A) and HCT116:PNUTS-AID (B) cell lines treated as in Figure 1 A fixed and subjected to FISH using pTRA-20 probe. Quantification of the area occupied by the FISH signals. Sample size: HCT116:RepoMan-AID: Control=317, IAA=262, HCT116:PNUTS-AID: Control=204, IAA=313.The box inside the violin represents the 75th and 25th percentile, whiskers indicate the upper and lower adjacent values and the line is the median. The data presented in the violin plots were statistically analysed with a Wilcoxon test. ns= not significant. E) Expression levels of CENPB obtained from the RNA seq experiments described in Figure 1 E. The values represent the average of the 3 independent replicas and the error bars are the standard deviations. The experiments were analysed by a Student’s t-test. *= p<0.05 F and G) Representative images of α-TUBULIN immunostaining using anti-α-TUBULIN antibody on HCT116:Ki-67-AID synchronised as indicated in Figure 1 A and treated without (F) or with

